# A Modular Framework for Automated Segmentation and Analysis of AFM Imaging of Chromatin Organization

**DOI:** 10.64898/2026.03.06.708946

**Authors:** Emily Winther Sørensen, Sushil Pangeni, Raquel Urteaga-Merino, Peter J. Murray, Sergei Rudnizky, Ting-Wei Liao, Fahad Rashid, Jihee Hwang, Maryam Yamadi, Xinyu A. Feng, Jonas Zähringer, Stephanie Gu, Iain F. Davidson, Laura Caccianini, Manuel Osorio-Valeriano, Lucas Farnung, Seychelle M. Vos, Jan-Michael Peters, James Berger, Carl Wu, Nikos S. Hatzakis, Julius B. Kirkegaard, Taekjip Ha

## Abstract

Chromatin organization underlies essential genome functions, but its nanoscale organization remains challenging to capture and quantify with precision. Atomic force microscopy (AFM) offers direct structural readouts of DNA and chromatin, yet translating these rich images into reproducible biological metrics has been limited by the lack of standardized, scalable analysis tools. Here we present DNAsight, an automated analysis framework that integrates machine learning (ML)-based segmentation with modular, base-pair-calibrated quantification of DNA spatial organization, looping, nucleosome spacing, and protein clustering. Applied across diverse chromatin-associated proteins, DNAsight reveals protein-specific organizational signatures, including topology-dependent compaction by integration host factor (IHF), condition-dependent changes in loop-like DNA structures in cohesin-CTCF-precocious dissociation of sisters 5A (PDS5A) reactions, and promoter-driven multimerization of GAGA factor (GAF) clusters. The framework further enables direct extraction of nucleosome spacing distributions from raw AFM images, providing a label-free route to investigate chromatin fiber architecture. Together, these advances establish DNAsight as a generalizable and scalable approach for converting AFM measurements into quantitative insights into the physical principles of chromatin organization.

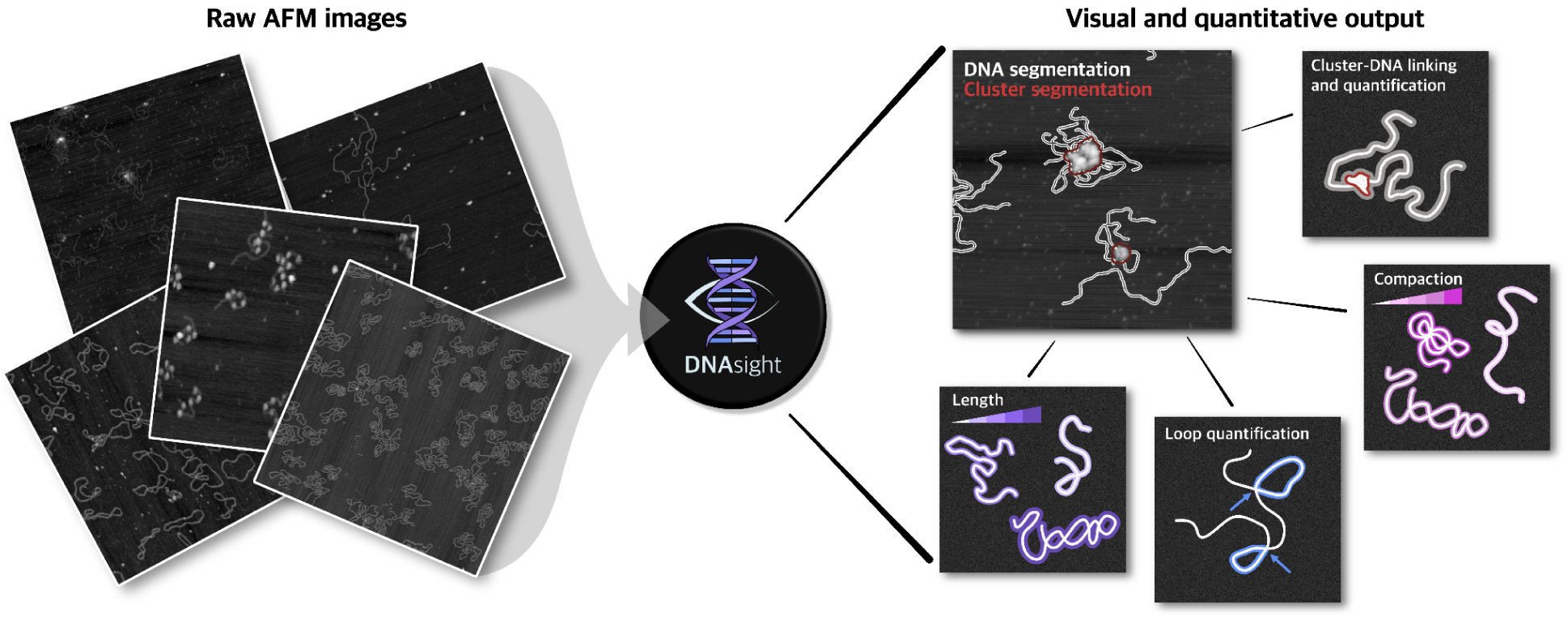

## INTRODUCTION

Understanding chromatin organization at the nanoscale is essential for deciphering the regulatory mechanisms that govern genome accessibility, gene expression, and nuclear architecture^1,2^. Here, atomic force microscopy (AFM) enables direct visualization of individual DNA-protein complexes under near-physiological conditions^3–5^, offering single-molecule (SM) resolution of DNA-protein organization. AFM imaging has revealed a broad spectrum of DNA and chromatin architectures, from naked and supercoiled DNA to nucleosome arrays, DNA-binding transcription factors, and higher-order chromatin assemblies, thus providing structural insights into chromatin topology across scales that help bridge the gap between atomic resolution structural studies and ensemble biochemical approaches^3,4,6,7^.

High-throughput AFM pipelines have been used to analyze nucleosome conformations^8^, allowing measurement of unwrapping states and height/shape differences across H3 vs. CENP-A nucleosomes^9^. AFM has also been used to characterize DNA compaction by polyamines^10^, revealing how multivalent cations such as spermidine or spermine can drive cooperative condensation of long DNA molecules into globular or toroidal structures^11^. Additionally, AFM has been used to explore how DNA-condensin interactions shape looping or bridging architectures^12,13^. Collectively, these studies demonstrate the versatility of AFM in resolving multiple organizational regimes of DNA, offering a comprehensive structural view that complements biochemical and genomic approaches.

Despite these strengths, extracting quantitative information from AFM images remains a key bottleneck. Many studies have relied on careful manual or semi-automated contour tracing to quantify DNA or chromatin features^9,14,15^. While such expert-guided approaches are reliable and can capture subtle structural details, they are also subjective and are cumbersome to scale to large datasets. Even when automated steps, such as background flattening, threshold-based segmentation, or particle detection, are introduced, these pipelines are often customized for a specific dataset and rarely generalize beyond their original context^16–18^.

Recent advances in computational AFM analysis have begun to address these limitations. Tools such as TopoStats^12,19^ have automated the tracing of DNA molecules from static AFM images, enabling reproducible measurement of contour length, curvature, and height profiles. Other image-analysis frameworks have applied classical computer-vision methods to extract DNA backbones or identify nucleosome positions^9,20,21^. However, these pipelines focus primarily on individual DNA molecules rather than complex chromatin architectures and often require substantial manual supervision during segmentation and feature selection. Nevertheless, these works have established an important precedent for bringing computational rigor to AFM studies of nucleic acids.

Building on this foundation, we introduce DNAsight, a modular computational framework that automates and standardizes the analysis of AFM images of chromatin. DNAsight integrates machine-learning (ML) based segmentation with quantification modules for DNA length, spatial organization, loop quantification, nucleosome spacing, and protein-mediated clustering. Beyond improving reproducibility and throughput, it expands the range of organizational features that can be systematically extracted from AFM data. The framework is designed to accommodate diverse experimental systems, here demonstrated on reconstituted chromatin fibers and chromatin-associated proteins such as GAGA factor (GAF), integration host factor (IHF), and cohesin regulators. Together, these results illustrate how automated analysis can broaden the scope of information extracted from AFM measurements of DNA and DNA-protein interactions, and provide new opportunities for quantitative studies of chromatin organization.

## METHODS AND MATERIALS

### Experimental methods

#### High-speed AFM

High Speed AFM, a commercial Sample-Scanning High-Speed Atomic Force Microscope (SS-NEX Ando model) from RIBM (Research Institute of Biomolecule Metrology Co., Ltd.), was used for experiments. Tapping mode was employed to minimize interference with the deposited sample, and all deposited samples were captured in solution. Ultra-Short Cantilevers (USC-F1.2k0.15-10), specifically designed for high-speed AFM with a resonance frequency of 1200 MHz, a spring constant of 0.15 N/m, and a length of 7 µm, were purchased from NanoAndMore and utilized in these experiments. A wide scanner was employed with scan speeds ranging from 0.05 to 6 second per frame, with the resolution set to 100×100 pixels to 500×500 pixels.

#### Surface passivations and DNA imaging

All experiments were carried out on mica functionalized with poly-lysine, closely following a published protocol^22^. Mica discs sized to fit the glass rod (and the corresponding rod) were obtained from RIBM. The mica disc was affixed to the glass rod with commercially available nail polish, and the rod was then installed on the AFM scanner. Immediately before each measurement, the mica was freshly cleaved by gently lifting the top layer with one-sided tape. The exposed surface was treated with poly(lysine) (MW ∼1000–5000; Sigma-Aldrich P0879) at 0.05 mg/mL for 3 min, followed by two rinses with distilled water. DNA prepared in the buffer of choice was diluted to 0.1-1 nM (as appropriate for each experiment), and 3 µL was applied to the poly-lysine-coated mica and allowed to adsorb for 3 min. The surface was then rinsed with distilled water and imaged in water using a high-speed AFM microscope. Although imaging in water may in principle perturb weakly bound or structurally unstable assemblies relative to their original buffer conditions, incomplete molecules were excluded during downstream filtering and analysis. AFM imaging of nucleic acids and nucleic acid–protein complexes in water has also been reported previously^23,24^.

#### AFM image conversions

HS-AFM images were viewed and analyzed using the software built by Prof. Toshio Ando’s laboratory-built software, Kodec 4.4.7.39, with available source code. Tilt and other image correction details are available in the literature^25^. Kodec was only used for the conversion of canonical asd files into bmp files that could be further used for the imaging analysis platforms discussed in the paper.

#### Preparation of supercoiled DNA substrates

Plasmid DNA (10 µg) was nicked with 35 U Nt.BbvCI in 210 µL 1× NEBuffer™ 2.1 (New England Biolabs). For low supercoiling, 400 U T4 DNA ligase and 5 U T4 DNA polymerase were added with 10 mM DTT and 2 mM ATP (final). For mid and high supercoiling, ethidium bromide (EtBr) was added to 4 µM or 10 µM (final), respectively. Reactions were incubated at 37 °C for 90 min, followed by treatment with 50 U T5 exonuclease at 37 °C for 60 min. DNA was purified using the QIAquick PCR Purification Kit (QIAGEN); EtBr-treated samples were subjected to butanol extraction prior to the final cleanup.

#### IHF purification

E. coli integration host factor (IHF; ihfA–ihfB operon cloned in pET21) was overexpressed in BL21-AI cells grown in 2×YT medium supplemented with carbenicillin at 37 °C. Expression was induced at an OD₆₀₀ of ∼0.8 with 0.5 mM IPTG and 2g/L arabinose for 3 h. Cells were harvested by centrifugation, resuspended in lysis buffer (25 mM Tris-HCl pH 7.4, 50 mM NaCl, 10% glycerol, 1 mM EDTA, 1 mM TCEP, and protease inhibitors), and lysed by incubation with lysozyme (1 mg mL⁻¹) on ice followed by sonication. Lysates were clarified by centrifugation at 18,000 rpm for 45 min. Nucleic acids were removed by dropwise addition of 8 M LiCl to a final concentration of 1 M at 4°C, followed by centrifugation. Proteins were fractionated by sequential ammonium sulfate precipitation, discarding the 50% saturation pellet and retaining the 80% saturation pellet. The resulting pellet was resuspended and dialyzed into high-salt HEPES buffer (20 mM HEPES pH 7.45, 400 mM NaCl, 10% glycerol, 1 mM EDTA, 1 mM TCEP), then purified by heparin affinity chromatography [Cytiva] using a 0.4–1.5 M NaCl gradient. Final polishing was performed by Superdex 75 size-exclusion chromatography (50 mM HEPES pH 7.5, 250 mM NaCl, 10% glycerol, 1 mM TCEP and 1 mM EDTA) [Cytiva]. Peak IHF fractions were pooled, aliquoted, flash-frozen in liquid nitrogen, and stored at −80 °C.

### GAF cluster measurements

GAF cluster experiments were performed on the passivated surface described above. GAF–DNA binding reactions were prepared as a 10× mixture containing 1 nM DNA and 10 nM GAF in GAF binding buffer (12.5 mM HEPES–KOH, pH 7.6, 0.05 mM EDTA, 6.25 mM MgCl₂, 50 mM NaCl) and incubated for 10 min. The reaction was then diluted 10-fold in the same buffer to 1× working conditions (final concentrations: 0.1 nM DNA and 1 nM GAF) to obtain an appropriate surface density, followed by an additional 10 min incubation. Samples were deposited onto the passivated surface and incubated for 10 min to allow adsorption. The surface was washed once with 80 µL ultrapure water, and imaging was performed in water.

During imaging, the setpoint/feedback parameters (and, when needed, the imaging height) were adjusted depending on cluster size to minimize tip–sample contact that can compress clusters or introduce blurring artifacts. GAF protein was expressed and purified as described previously^26^.

### IHF measurements

IHF cluster experiments were performed on the same passivated surface and followed the same surface deposition, wash, and imaging procedure described above (including adjustment of imaging parameters for large assemblies to avoid tip-induced artifacts). IHF–DNA binding reactions were prepared as a 10× mixture containing 1 nM DNA and 10 nM IHF in buffer (50 mM HEPES, pH 7.5, 100 mM NaCl) and incubated for 10 min. The reaction was then diluted 10-fold to 1× working conditions (final concentrations: 0.1 nM DNA and 1 nM IHF) to achieve an appropriate surface density, followed by an additional 10 min incubation. Samples were then deposited onto the passivated surface, incubated for 10 min, washed once with 80 µL ultrapure water, and imaged in water.

### Cohesin-CTCF-PDS5A looping measurements

A 7,979 bp DNA substrate containing two CTCF consensus motifs, the center of which were 511 or 514 bp from each DNA end), cohesin, STAG1, NIPBL-MAU2 (“NIPBL”), PDS5A, and CTCF were prepared as previously described^27^. Cohesin and STAG1 were preincubated each at 25 nM in SMC buffer (50 mM HEPES pH 7.5, 100 mM NaCl, 10 mM MgCl_2_) for 10 minutes on ice. The cohesin reaction was performed by mixing 0.5 nM DNA, 5 nM CTCF, 7.5 nM cohesin-STAG1, 5 nM NIPBL, and 5 mM ATP in SMC buffer and incubating at 37°C for 5 minutes. For the PDS5A condition, 5 nM PDS5A was added after the 5 minutes and a subsequent 1 minute at 37°C incubation was performed. Following the incubation, the sample was diluted 5x in SMC buffer supplemented with 5 mM ATP and immediately deposited on a P-lysine-coated mica surface. The sample was incubated for 10 minutes on the surface, washed with 80 uL of water then imaged in water. The utilized proteins were expressed and purified as explained in a recent preprint^27^.

### Nucleosome reconstitutions

*Nucleosomal DNA construct formation:* Plasmids containing two or six repeats of the 601 nucleosome positioning sequence, separated by 80 bps of linker DNA, were cut using restriction enzymes to generate di and hexanucleosomal DNA (394 and 1364bps, respectively). The DNA fragments contained an additional 64 and 20bps linker on the hexanucleosomal sequence, and 20 bps at one end of the dinucleosome. The following restriction enzymes were used: BstZ17-HF and XhoI - dinucleosomes, and PciI and XhoI - hexanucleosomes. To generate non-well positioned nucleosomes, the ScaI enzyme was used to generate a single cut on a 4kb plasmid. Digested DNA was ethanol precipitated, and PAGE purified using Bio-Rad Prep cells of different acrylamide/bisacrylamide percentage.

*Reconstitution of polynucleosomes:* Nucleosomes were reconstituted using salt gradient dialysis. Human histone octamer was titrated at around 6X, 2X or 5X molar fold excess to DNA concentration depending on the nucleosome construct (hexa-, di-or non-well positioned, respectively). BSA was not included in reconstitutions for cleaner AFM nucleosome signals. Efficient assembly was checked on a native PAGE gel (3.5% hexa and non-well positioned, 6% dinucleosomes). Reactions were pulled down and purified over a 15-40% sucrose gradient (SW60Ti rotor, 45,000 rpm, 16hrs, 4C). Peak fractions were pooled, concentrated and buffer exchanged to nucleosome storage buffer (10mM Tris pH7.5, 1mM EDTA, 50mM NaCl, 3.5mM BME, 0.02% NP40).

### Nucleosome imaging

Nucleosomes were reconstituted as explained in the reconstitution section. A low salt buffer (LSB) was used to image the nucleosome. LSB buffer is 10 mM Tris pH 7.5 50 mM NaCl 1 mM EDTA 5mM BME 0.02% NP40. Briefly, nucleosomes were diluted to equivalent of 0.1 nM in concentration in LSB and deposited to the poly-lysine coated mica surface. Note that a low salt buffer is essential to provide the integrity of nucleosomal constructs. Then 3 uL was deposited on the surface and incubated for 5 minutes. Then samples were washed with LSB buffer once then imaged in water in the reservoir. Scan size was varied between 2000 nm*2000 nm to 300 nm*300 nm based on the size of the constructs.

### DNAsight

#### DNA segmentation

*Manual annotation:* AFM image output was converted into TIFF format (or other ImageJ compatible format like BMP). Hereafter, the images were annotated using the “Freehand Line” tool in ImageJ and the files were saved in TIFF format with the manual annotations. For this work, 258 images were manually annotated.

*Data acquisition and pre-processing:* TIFF images containing raw AFM data and corresponding manual annotation masks embedded as metadata (ImageJ overlays) were used to create a training dataset for DNA segmentation. Images were normalized by scaling pixel intensities to a range between 0 and 1, ensuring consistent contrast and brightness levels across all samples. This normalization provided uniform input data, reducing potential bias due to variable imaging conditions.

*Data augmentation:* To expand the effective dataset and improve model generalization, a diverse set of image augmentations was applied. Each image could be randomly rotated by up to ±30° and cropped to remove borders, while random horizontal and vertical flips introduced orientation variability. Moderate changes in brightness and contrast (±10%) simulated variations in imaging conditions. Additional stochastic perturbations included either image downscaling (to 30–90% of original resolution) or advanced blurring, mimicking differences in AFM tip sharpness or focus. Gaussian noise (σ = 0.01–0.05) and sharpening were also added to emulate sensor noise and enhance structural variability. Finally, all images were randomly cropped to 128 × 128 pixels.

*Generation of proximity maps:* Rather than using binary annotations as ground truth, the model was trained using continuous backbone-proximity maps derived from manual DNA annotations. These maps were generated by computing the Euclidean distance to the nearest annotated DNA contour for each pixel. Distances were then clamped to a predefined maximum (default: 5 pixels) to limit the spatial extent of the signal. The clamped distance values were subsequently inverted and normalized to the range [0, 1], such that pixels on the DNA backbone attain the maximum value and values decay smoothly with increasing distance into the background. This transformation yields a gradient-rich target representation that emphasizes the DNA medial axis and nearby boundary regions, providing a more informative and stable supervision signal during training.

*Model architecture:* The segmentation was performed using a convolutional neural network based on the widely established U-Net architecture. This model consists of two main pathways: (1) an encoder pathway (contracting path) that captures context by progressively reducing spatial dimensions through convolutional and max-pooling operations, and (2) a decoder pathway (expanding path) that restores spatial resolution through upsampling and convolutional layers. Skip connections between the encoder and decoder pathways ensure the preservation of detailed spatial information, critical for accurately delineating narrow DNA strands. The final layer of the model employs a 1×1 convolution followed by a sigmoid activation function, outputting a predicted normalized proximity map with pixel values between 0 and 1.

*Training procedure:* The U-Net model was trained using an Adam optimizer with a learning rate of 0.001 over multiple epochs (default 1500 epochs). During each iteration, batches of augmented images, their corresponding binary annotation masks, and pre-computed proximity maps were input into the model. Model weights were updated through gradient descent optimization to minimize prediction error.

*Loss function:* Training employed a customized WMSE loss function designed specifically for proximity map regression:

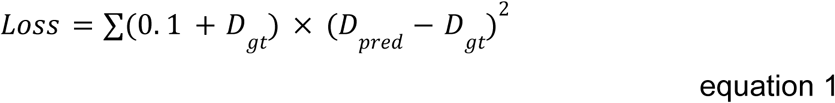

Here, 𝐷𝑔𝑡 represents the ground truth proximity map, and 𝐷𝑝𝑟𝑒𝑑 is the predicted map.

The weighting term 0.1+𝐷𝑔𝑡 ensures that errors near the boundaries – where 𝐷𝑔𝑡 is higher - are penalized more heavily, thus encouraging the network to focus on these critical regions. This approach provides a more refined supervisory signal compared to a standard binary segmentation loss and helps the model learn the fine details essential for accurate segmentation of DNA structures.

*Prediction and post-processing:* After the AFM images were passed through the trained network to produce continuous proximity maps, these were converted into skeletonized DNA backbones through a multi-step process. First, a high-confidence mask was defined by thresholding the proximity map. Second, small objects below a minimum area were removed to eliminate spurious detections. Third, backbone ridges were enhanced by a DoG filter, and a binary segmentation was created by combining this ridge response with the high-confidence proximity map. This binary segmentation was reduced to a one-pixel-wide backbone using skeletonization, and the skeleton was restricted to the previously validated components to ensure continuity within true DNA molecules. Finally, each skeletonized molecule was assigned a globally unique identifier, ensuring consistent indexing across all images in a dataset. Processed outputs were saved as two-layer image files: one layer containing the raw AFM image, the other containing the global skeleton ID map for the input folder. These global IDs are used and saved for all downstream analysis.

*Performance:* Segmentation accuracy was assessed using pixel-distance error between predicted skeletons and manually annotated centerlines. Akin to dynamic time warping^28,29^, we calculate the average of the minimum distance from prediction to label and label to prediction (see Figure S1). For assessing performance, a separate model was trained on 90% (232 images) of the data randomly chosen, and then evaluated on the remaining 10% (26 images) of the data.

### DNA length calibration and pixel-to-bp conversion

*DNA backbone length measurement from skeletons:* DNA lengths were measured directly from skeletonized backbones generated by the segmentation pipeline. Each skeletonized component was represented as a graph on the pixel grid with full eight-neighbour connectivity, allowing both orthogonal and diagonal steps. To avoid artificially shortened paths at branch points, diagonal “corner cuts” across square junctions were excluded. The skeleton graph was then decomposed into continuous path segments by tracing along edges until reaching either an endpoint or a junction. For closed contours where all nodes had degree two, the entire loop was returned as a single segment. The Euclidean length of each segment was computed as the sum of one unit for orthogonal steps and √2 for diagonal steps, and the lengths of all segments belonging to a component were added to obtain its total contour length in pixels. As the length was calculated from the center of the pixels, half the length of each end-point pixel was added to the length (√2/2 for diagonal pixels and ½ for straight). Components that touched the image boundary were flagged as edge-touching to indicate possible partial molecules.

*Calibration constant by filtering:* For calibration, we used reference DNA molecules of known sequence length. To improve robustness of the calibration in datasets containing truncated molecules, overlapping DNA, edge-touching molecules, impurities, or occasional segmentation artifacts, DNAsight allows optional filtering of the reference population before estimating the calibration constant. In the implementation used here, edge-touching molecules were excluded and additional quantile-based filtering was applied using user-defined lower and upper bounds (default: 25th and 75th percentiles).

These thresholds are not fixed requirements of the framework and can be relaxed or disabled depending on dataset quality. For highly pure, well-resolved reference datasets, the full distribution can in principle be retained for calibration. The purpose of this filtering step is to obtain a stable estimate of the calibration constant from intact single reference molecules, rather than to characterize the full spread of the raw sample.

This filtering step removed atypical fragments, overlapping molecules, and rare segmentation artifacts. After filtering, pixel lengths were converted to nanometers by multiplying by the pixel size recorded from the AFM scan settings. The calibration constant, expressed as nanometers per bp, was then estimated as the mean length in nanometers of the retained molecules divided by the known sequence length in bps (equation 2).

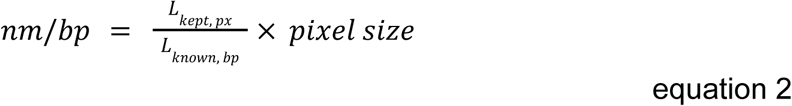

Where 𝐿𝑘𝑒𝑝𝑡, 𝑝𝑥 is the mean length of the kept DNA molecules after filtration in pixels and 𝐿𝑘𝑛𝑜𝑤𝑛, 𝑏𝑝 is the known length of the DNA molecules in bp. The spread of values across the filtered set provided a measure of variation and reproducibility of the calibration. Using a calibration

### Geometric features

*Compaction:* For each labeled DNA molecule, we quantify spatial spread using the radius of gyration 𝑅_𝑔_. Let {𝑟*_i_*}*^N^*𝑖=1 be the pixel coordinates of all pixels belonging to that DNA. We first compute the geometric centroid (center of mass of the pixel set) as

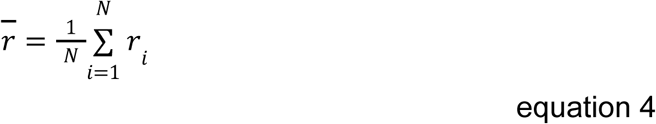

Hereafter the root-mean-square distance from the center is calculated as

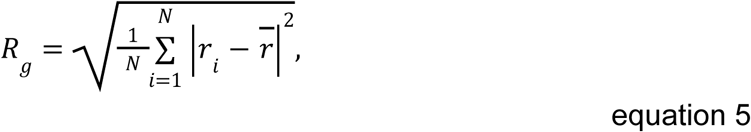

Intuitively, 𝑅𝑔 is small when the DNA pixels are tightly clustered around the centroid (compact object) and larger when pixels are spread out. Because 𝑅_𝑔_ naturally grows with the size of the object, we report the normalized and rescaled compaction, C, as seen in equation 6, where 𝐿 is the total skeleton length of that DNA in pixels. Multiplying by 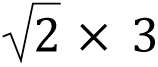 and subtracting from 1 rescales the compaction to be between 0 and 1, for easier interpretability.

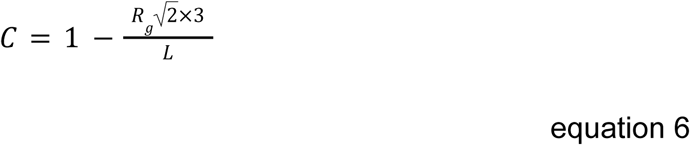

This normalization makes values comparable across molecules of different lengths.

However, not across different pixel sizes. As such, when pixel size is provided 𝐶 is further normalized to be comparable across varying pixel sizes.

*Crossings:* To find branching points or crosses, we identify pixels along the DNA mask which have ≥ 3 neighbouring annotated pixels in the DNA mask. Hereafter, nearby junctions are merged within a 3 pixel radius to not overestimate the number of crossings.

*Tortuiosity:* We define tortuosity as the ratio of path length to the most distant end-to-end chord measured on the skeleton (equation 7).

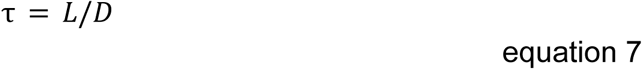

Where 𝐿 is the total skeleton length (pixels) and 𝐷 is the Euclidean distance between two endpoints on the skeleton. In the case of >2 endpoints in the skeleton, the two points furthest from each other are used. For closed loops (no endpoints), τ is undefined and omitted.

*Curvature:* Obtaining a single representative centerline per molecule is generally not possible, and when it is, finding such a path is computational infeasible (NP hard). As a compromise, we traverse the graph by favoring straight continuation across junctions (as defined in crossings). Starting from an endpoint (or a crossing if no endpoints exist), we repeatedly choose the outgoing edge whose direction best aligns with the current travel direction. Direction is estimated from the last 𝑘 steps (look-ahead/behind, 𝑘 = 3 by default), and alignment is measured by the dot product (i.e., the cosine of the turning angle). The traversal proceeds until no unused edges remain along a straight continuation. Among all such traversals, we retain the longest resulting path (maximum arclength in pixels). For closed loops (no endpoints), we walk the contours to produce the longest simple path available. This strategy allows us to follow the biologically most likely path when extracting curvature

The selected path (ordered pixel coordinates (𝑥𝑖, 𝑦𝑖)) is lightly smoothed with a Savitzky–Golay filter to reduce pixel anisotropy and discretization noise introduced by skeletonization while preserving local geometry. Because this smoothing can influence absolute curvature values, the smoothing window is a user-adjustable parameter rather than a fixed requirement of the framework (note that it must be an odd number). Coordinates are then re-sampled to approximately uniform arclength spacing Δ𝑠 (default ≈ 1 px).

We hereafter fit natural cubic splines 𝑥(𝑠) and 𝑦(𝑠) to the arclength-parameterized path and compute the unsigned curvature,

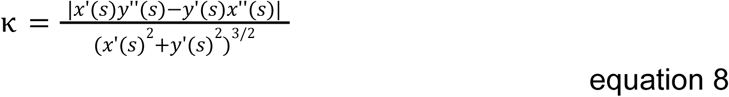

To mitigate end effects, we discard a small fraction of points at both ends (default 2%). When pixel-to-nm calibration is available, κ in 𝑝𝑥^−1^ is converted to 𝑛𝑚^−1^ by dividing by pixel size.

Per molecule, we report the mean, std, minimum and maximum curvature computed over the trimmed spline sample. In the analyses presented here, we used a default smoothing window of 15 pixels. Sensitivity analysis across smoothing-window sizes showed that very small windows produced higher and more variable curvature estimates, consistent with stronger sensitivity to pixel-scale jaggedness, whereas the estimates stabilized over at window sizes around 11-17 (Figure S2). Paths shorter than four points after preprocessing are excluded. If smoothing parameters are too large for short segments, the window is adaptively reduced to the nearest valid odd value (minimum 𝑤 = 3) to ensure a stable fit.

*Strong bends:* We detect strong bends from the unsigned curvature profile |κ(𝑠)| sampled at approximately uniform arc-length positions. Using a θ𝑚𝑖𝑛 = 30 degrees and 𝑙𝑚𝑖𝑛 = 3 𝑝𝑥, we set a local curvature threshold as defined in equation 9 and a maintenance threshold κ_𝑘𝑒𝑒𝑝_ = 0. 9 × κ_𝑡ℎ𝑟_.

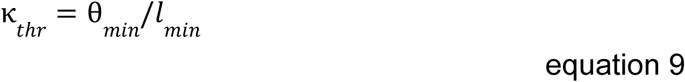

A bend begins at the first sample where |κ| ≥ 𝑘_𝑡ℎ𝑟_ and continuous while |κ| ≥ κ_𝑘𝑒𝑒𝑝_; any dip longer than 1 px below κ_𝑘𝑒𝑒𝑝_ immediately ends the run. If the accumulated span of the run reaches at least 𝑙𝑚𝑖𝑛, the event is counted as one strong bend; otherwise it is discarded. Counting then resumes after the end of the run so that events do not overlap. Curvature is in 𝑝𝑥^−1^ (or 𝑛𝑚^−1^ if upstream pixel-size calibration is applied).

*Elongation:* Elongation is derived from the best-fit ellipse to each connected DNA mask. We report elongation as calculated as the ratio of the major to minor axis lengths (𝑎, 𝑏) from region properties

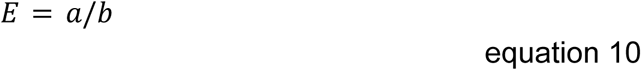

Larger 𝐸 indicates more anisotropic rod-like shapes.

### Loop detection and quantification

For each segmented DNA, we searched for interior holes in the complement of the DNA mask (background) using connected-component labeling with border clearing, so that only fully enclosed holes were retained. A hole was considered a loop candidate if its contact length was above a minimum threshold (default ≥ 10 pixels).

To represent each candidate loop by a single curve, we skeletonized the contacting-pixel set to one-pixel width and suppressed spurious branching. Branch points on the skeleton were detected by hit-or-miss transforms using a bank of rotated/reflected 3×3 templates, then pruned and relabeled to isolate the largest contacting strand. Branch points were then restored and the result re-skeletonized to yield a clean one-pixel curve along the DNA-hole interface.

To remove pseudo loops on the same DNA (enclosed structures with many associated loops), we additionally filtered strongly overlapping loop candidates. Specifically, when two candidate loop paths overlapped by at least 5 pixels, candidates were sorted by loop size and the smaller candidate was preferentially retained. The rationale for this choice is that the larger enclosed contour often arose when a longer DNA segment bent back around an existing loop and re-contacted the DNA along its side, thereby generating an additional apparent closed contour in the 2D segmentation rather than a second independent looping event. Such geometries generally require a relatively long contour segment to extend away from the primary loop and return into contact. In this sense, the larger candidate is often more plausibly explained as a secondary enclosure created by self-overlap of an extended DNA segment than as a distinct loop. Retaining the smaller candidate was therefore used as a pragmatic way to reduce false-positive loop calls, while acknowledging that this distinction is not exact in all cases.

For each retained loop, we are interested in identifying the loop location, which we define as the shortest distance of the loop attachment point to either DNA end. This nearest-end metric is convenient for comparing loops across molecules, but it as blind to the directionality of linear DNA as the measurement is. Consequently, nonspecific background loops may fall by chance within a site-proximal window and be scored as apparent site-associated events, creating a risk of false positives. The metric is therefore best suited to symmetric constructs, where equivalent sites are located at the same distance from either end. However, in either case, because total DNA length is also measured, the complementary distance to the opposite end can additionally be calculated to inspect the full positional distribution. Together, these descriptors are therefore most useful for identifying positional trends or enrichments at the population level, rather than for making absolute assignments of individual loop-like structures to specific anchoring events.

We identified the attachment point on the loop by removing the loop from the DNA mask, mildly smoothing the residual mask, and selecting the loop pixel with maximal contact to the remaining DNA; this point was then snapped to the nearest skeleton pixel on the DNA centerline. We then computed distances from this attachment to each true DNA endpoint (as explained in the DNA length calculation section), and reported the shortest of these. This distance includes loops along the path.

Note that distance to the nearest end can be longer than half the length of the DNA in the case of the DNA having only one end point (the other end going over the edge of the image) or if the two ends of the DNA segmentation branch from the same single line skeleton DNA.

### Cluster segmentation

To allow for segmentation of a wide range of sizes and shapes of clusters, we created two options of cluster segmentation. One for larger assemblies of various shapes and one for smaller circular clusters. Both segmentation types have the same output, where a global ID is saved and used for all downstream quantifications.

*Large assemblies:* AFM frames were segmented to isolate large protein assemblies with a marker-based random-walker approach^30^. Images were lightly smoothed (Gaussian, σ = 1 𝑝𝑥) to suppress pixel noise. A coarse foreground seed was obtained by thresholding the blurred image at the desired multiplication above the mean intensity of the whole image. To provide robust markers for the random-walker, the seed was dilated to obtain confident foreground regions, and its complement was dilated to obtain confident background regions (both dilation parameters are user inputs).

A binary random-walker diffusion assigned each pixel to foreground or background given these markers. Small interior voids were filled, and connected components were extracted with 8-connectivity. Components with an area below a minimum size (user input) were discarded. For each retained cluster we recorded: centroid (center of mass), area (pixel count), and integrated intensity (sum of grayscale values within the region). To enable dataset-level analyses, clusters across all images were assigned global IDs and saved alongside per-image overlays.

*Small clusters:* Small puncta were handled by a two-stage “detect-then-segment” pipeline optimized for near-round spots. Firstly, candidate spots were first localized using a classical particle detection method^31,32^, using an expected spot diameter (user input), separation at least as large as the diameter, and an intensity/mass cutoff to suppress noise. Around each center we cropped a small window (user input) and produced a binary mask by either Otsu thresholding or a fixed percentile threshold. Very small specks were removed. If multiple blobs appeared in the window, the blob containing (or nearest to) the detection center was kept; otherwise a small fallback disk was used to represent the spot at image borders.

Because windows can overlap, the stack of per-spot masks was converted into a single exclusive label map by assigning any overlapping pixel to the nearest spot center (nearest-center rule). Labeled spots were then screened with simple geometric criteria. We required the area to fall within a practical range (user input). Shape quality was assessed by circularity 𝐶=4π𝐴/𝑃^2^, eccentricity, and solidity. For each spot that passed screening, we recorded the centroid, area, and integrated intensity, and assigned a dataset-wide global identifier. Outputs share the same schema as the large-assembly pipeline.

### Cluster quantification and background correction

Following segmentation, each detected protein cluster was quantified for its integrated intensity and projected area, with per-image normalization and local background subtraction to allow comparison across samples and imaging conditions.

For each cluster, the segmentation mask defines the pixel region corresponding to that cluster. The summed intensity (sum of pixel values within the mask) and the mask area in pixels were extracted directly from the original AFM or fluorescence image.

To correct for the local background, a ring-based method was used. Two binary dilations of the cluster mask were generated - one expanding the cluster boundary by 2 pixels and another by 5 pixels. The ring-shaped region between these two dilations represents a local shell surrounding the cluster but excluding the cluster itself. The mean pixel intensity within this ring was taken as the local background intensity. The background-corrected integrated intensity for each cluster was then computed as

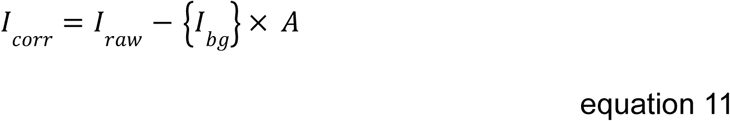

Where 𝐼𝑟𝑎𝑤 is the raw summed intensity inside the cluster, {𝐼𝑏𝑔} is the average intensity in the background ring, and 𝐴 is the cluster area in pixels. For each cluster, additional parameters were derived; cluster area (px, nm²) - number of pixels multiplied by the squared pixel size; summed intensity per nm or per nm² - intensity normalized by area or linear scale; background-corrected intensities - both absolute and normalized forms as above; and edge-touch flag - a Boolean flag identifying clusters that intersected the image border, to allow optional exclusion from quantitative analyses. All corrected and normalized values were exported to a summary table (see supporting information for details).

### Cluster-DNA linking and quantification

*Linking segmentations:* To quantitatively relate DNA structures to the identified GAF clusters, a region-growing strategy was implemented. For each cluster, its binary mask was expanded using morphological dilation to account for positional uncertainty. This expanded cluster mask was then used to “capture” adjacent DNA components from the binary DNA segmentation.

### Analytical methods

#### SNR of DNA molecules and images

Manual DNA annotations were used as primary segmentation masks. To define measurement regions, three concentric zones were constructed by morphological dilation of the segmentation mask.

1. **Signal region:** the segmentation mask dilated outward by 1 pixel, capturing the DNA signal while remaining robust to slight under-annotation.
2. **Gap ring (ignored):** a 1-pixel-wide buffer immediately outside the signal region (between +1 px and +2 px dilations), excluded to prevent signal bleed or partial-volume effects from contaminating the background estimate.
3. **Background ring:** a 2-pixel-wide annulus immediately outside the gap (between+2 px and +4 px dilations), providing a local background matched to the same imaging field and surface.

For each image, signal (µ𝑠) was defined as the mean intensity of all signal-region pixels across all DNA molecules identified by the segmentation mask. Background mean (µ_𝑏_) and background standard deviation (σ_𝑏_) were defined analogously from the background rings pooled across the same segmentation mask.

The image-level SNR was defined as

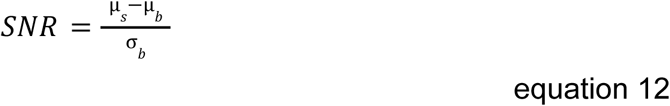

Pooling signal and background statistics across the segmentation mask yields a single, robust SNR per image that is insensitive to local intensity variations along individual DNA molecules. The ignored gap prevents residual DNA signal from inflating background estimates, while the local background annulus controls for slow spatial variations in the AFM surface and scanner.

### Dominant peak and fit window for DNA contour length estimation

To estimate the characteristic DNA length while minimizing contributions from incomplete or overlapping molecules, a Gaussian model was fitted only to the dominant population in the length distribution. Here, a histogram of DNA contour lengths was constructed using a fixed number of bins. The center of the histogram bin with the highest occupancy was identified and taken as an initial estimate of the dominant peak position. To determine an initial width of the distribution in a manner robust to outliers, the median absolute deviation (MAD) of the full length distribution was calculated and converted to an equivalent Gaussian standard deviation using a factor of 1.4826^33^. This value was used to define an initial fitting window centered on the dominant peak.

All DNA molecules with contour lengths within ±2.5 times the estimated standard deviation around the peak center were included in the fit. A Gaussian distribution was then fitted to the selected subset using maximum likelihood estimation, yielding the mean contour length and standard deviation reported for the dataset.

### Fitting of lognormal mixture model to nucleosome-associated DNA segment lengths

We estimated linker-length heterogeneity of the poly-nucleosomes by fitting a two-component lognormal mixture to the distribution of measured lengths of nucleosome-associated DNA segments. Before modeling, we removed groups whose constituent DNA contours touched the image boundary. The remaining per-group linker lists were flattened to a single sample of lengths, then restricted to positive values up to 500 bp to focus the model on the primary modes and avoid extreme tails. We then applied a Gaussian mixture model to the natural-log of the lengths and used AIC and BIC to determine the ideal number of states to be two. Model parameters were estimated by maximum likelihood via the expectation-maximization algorithm, allowing full covariance for each component. The fitted means and variances in log space and the mixing proportions were then mapped back to linear units to report the mean.

## RESULTS

### DNAsight: A Modular Framework for Analysis of Chromatin in AFM Images

To systematically extract structural information from DNA and chromatin imaged by AFM, we developed DNAsight, a modular image-analysis framework that links segmentation with biologically meaningful quantification (Figure 1). The pipeline is organized into two segmentation modules for DNA and proteins/clusters (Figure 1A) and four quantification modules (Figure 1B). The modules are designed to work both sequentially and independently, with some modules being coupled, depending on the experimental question. As input, DNAsight operates on raw AFM images of DNA or DNA-protein complexes and outputs standardized, interpretable metrics, such as DNA length, spatial organization, looping and DNA-protein clustering.

**Figure 1.**
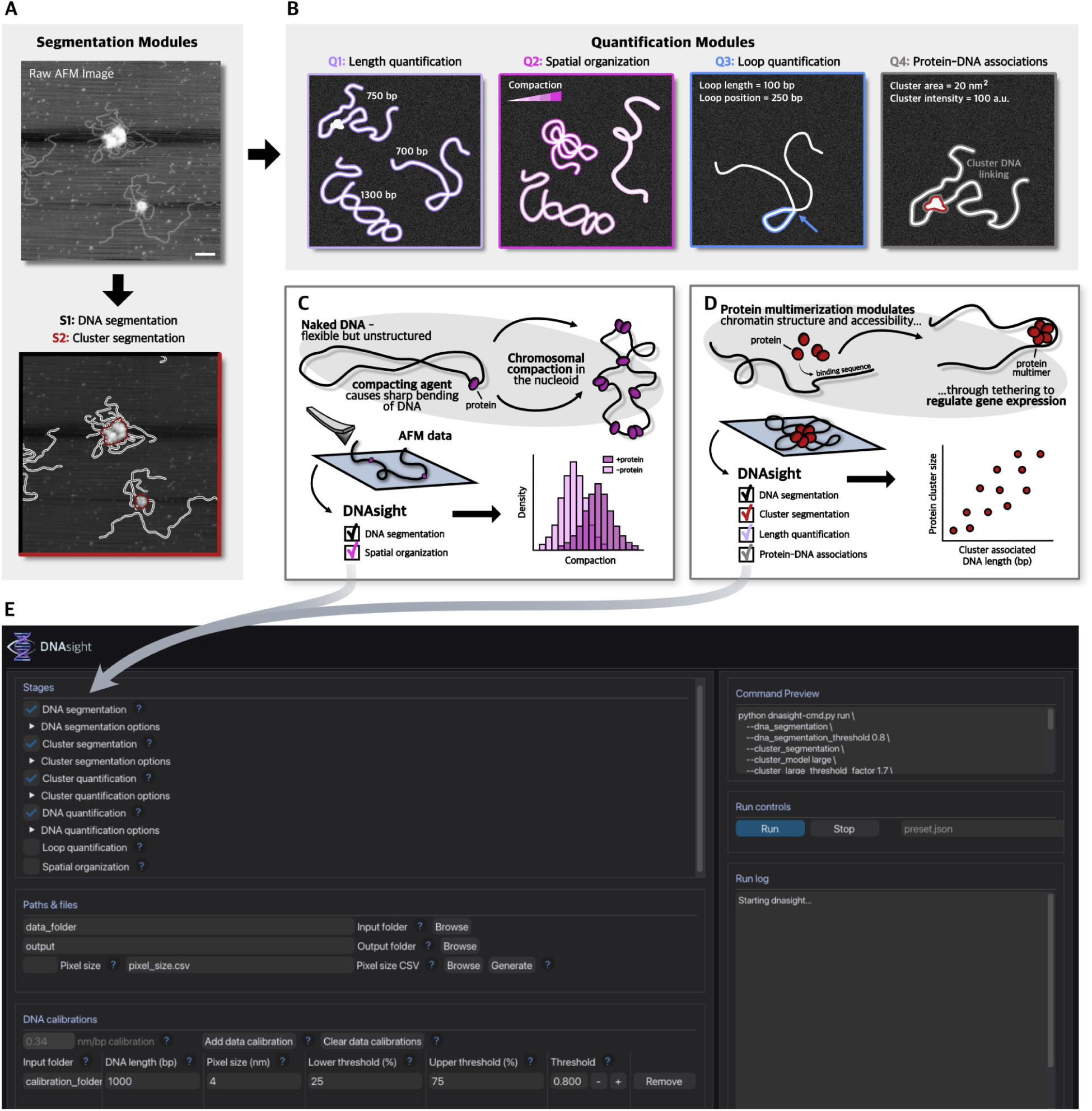
A schematic representation of DNAsight contents and pipeline. (A) Segmentation modules, demonstrated on AFM image, S1: DNA segmentation (white outline) and S2: Cluster segmentation (red outline). Shown applied to an AFM image of GAF proteins and DNA (scale bar 100 nm). (B) Cartoon representations of post-segmentations quantification modules Q1: DNA length quantification, Q2: DNA spatial organization, Q3: DNA loop quantification and Q4: Protein-DNA associations. (C) Possible mock application of DNAsight using S1 and Q2 to quantify DNA compaction caused by a compacting agent protein, real example shown later. (D) Another possible mock application of DNAsight using S1, S2, Q1 and Q4 to quantify clustering of a multimerizing protein and its correlation with DNA lengths, real example shown later. (E) Screenshot of the free-to-use GUI that integrates the multimodular pipeline of DNAsight. The user-friendly GUI allows users to perform all core functions of DNAsight directly on raw AFM images, both including segmentation and quantifications, as well as plotting and evaluation of data.

The DNA segmentation module (S1) employs an ML-based approach to trace DNA molecules, establishing the structural skeleton for downstream analyses. The protein segmentation module (S2) detects protein features and clusters across multiple size scales, from compact puncta such as nucleosome-like particles to larger protein assemblies, using intensity-based thresholding and morphological filtering tailored to each regime. Downstream linking enables integration of protein architecture with the underlying DNA landscape. Together, S1 and S2 define the structural basis for all subsequent quantifications.

The subsequent quantification modules (Q1-Q4) convert these segmentations into biological descriptors (Figure 1B-D). Q1 determines DNA lengths in nanometers and base-pairs (bps) using reference molecules of known size, ensuring that all subsequent measurements are biologically interpretable and tailored to the experimental system. Q2 quantifies geometric features of DNA, including measures of compaction, curvature, tortuosity, and shape, thereby allowing systematic comparison of chromatin states across conditions. Q3 uses skeletonized DNA masks from S1 to detect loop-like structures, reporting their positions and lengths relative to DNA ends. Here, loops, are defined as regions where the DNA backbone folds back and approaches or crosses itself, forming a topologically distinct closed or near-closed segment. Finally, Q4 integrates the DNA and protein masks generated in S1 and S2 to quantify the spatial relationship between proteins and DNA. This includes identifying where protein complexes bind along individual DNA molecules, measuring distances between adjacent binding sites or proteins as well as determining cluster statistics in relation to DNA length or position. The modularity of DNAsight allows users to select relevant analysis modules to their experimental question (Figure 1C–D). For instance, combining S1 with Q2 enables quantification of DNA compaction induced by a compacting agent (Figure 1C), while integrating S1, S2, Q1, and Q4 allows measurement of protein clustering and its correlation with the length of recruited DNA (Figure 1D).

DNAsight is implemented as a modular framework that allows users to select analysis modules based on the experimental question. A graphical user interface (GUI) provides guided access from raw AFM images to quantitative outputs (Figure 1E), while detailed usage instructions are provided in the online documentation available through the GitHub repository.

### Automated DNA Tracing Achieves Human-Level Precision and Base-Pair Calibration Across Imaging Conditions

At the core of DNAsight is the DNA segmentation module (S1), which extracts SM DNA segmentations directly from AFM topography images. Rather than predicting a binary DNA mask, we trained a U-Net convolutional neural network to output continuous backbone-proximity maps, where each pixel encodes its normalized proximity to the nearest DNA centerline. In these maps, values are maximal on the DNA backbone and decay smoothly toward zero with increasing distance into the background. This representation addresses the extreme class imbalance in AFM images, where DNA typically occupies only ∼1% of pixels. By spreading DNA information into the surrounding background, proximity maps ensure that every pixel contributes to the learning process. To further counteract imbalance, we optimized the model with a customized weighted mean squared error (WMSE) loss, in which pixels close to the backbone contribute most strongly while background pixels retain a small baseline weight (Methods and Materials). Together, the proximity map formulation and WMSE loss prevent trivial all-background predictions, while prioritizing accurate recovery of the DNA medial axis (Figure 2A).

**Figure 2.**
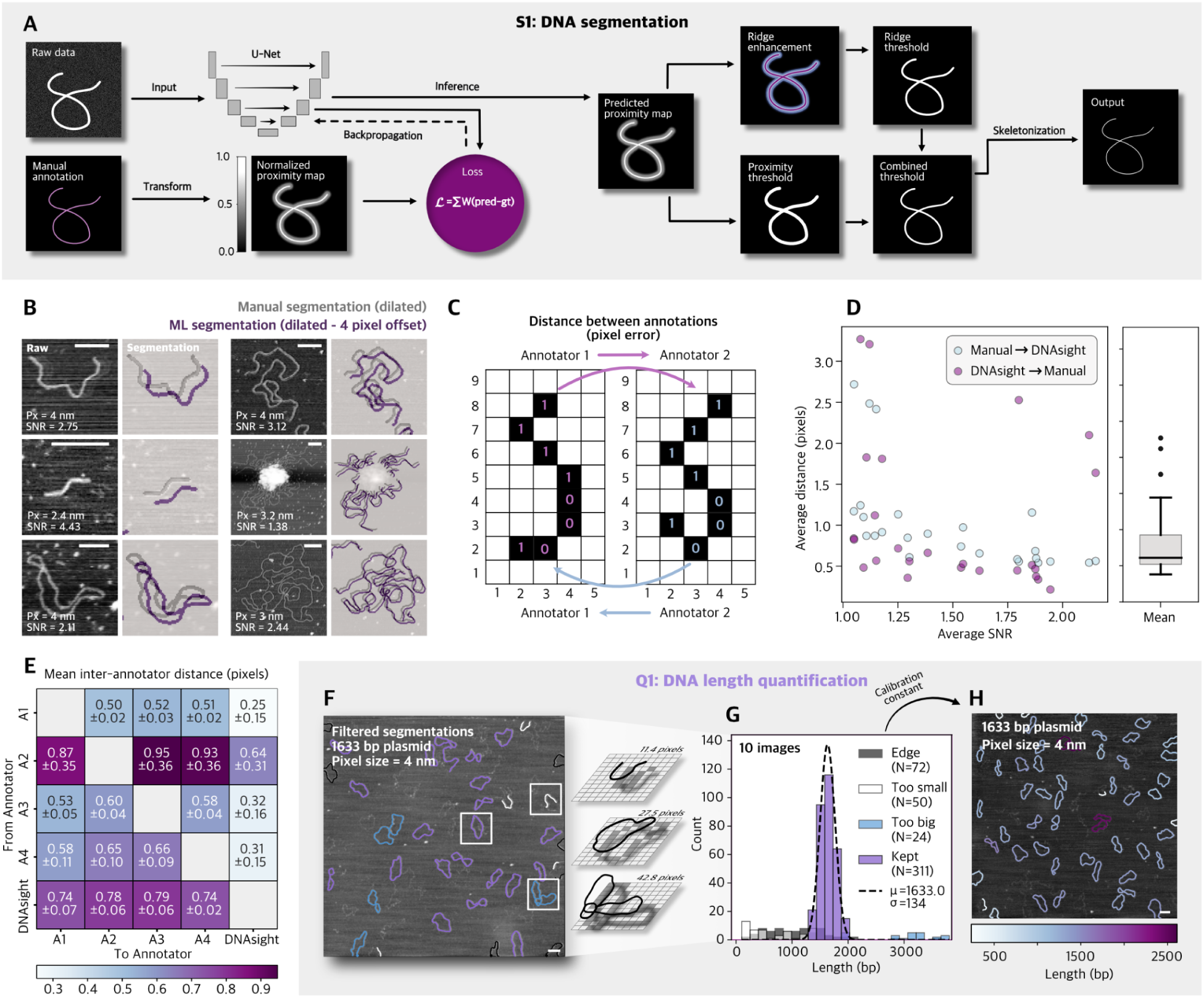
DNA segmentation and conversion to bp. (A) Workflow of the DNA segmentation module (S1). AFM topographs are processed by a U-Net trained on normalized proximity maps derived from manual annotations. Predicted proximity maps are refined through watershed segmentation, area filtering, and a DoG (ridge enhancement) filter to produce one-pixel-wide skeletons suitable for downstream analysis. (B) DNA segmentation results (purple) compared to manual annotation (grey) across different pixel sizes (Px) and SNRs (scale bar 100 nm). (C) Cartoon outline of inter-annotator pixel error measure. (D) Segmentation accuracy compared against a single manual annotation across SNRs, evaluated using pixel-distance error, shown in both directions, from DNAsight to manual (purple) and manual to DNAsight (blue) and the average (grey boxplot). (E) Pixel-distance error comparing 4 independent human annotators with DNAsight segmentation. (F) DNA length quantification module (Q1). Known-length molecules, here 1633 bp plasmids and linear fragments, are filtered post segmentation to exclude outlier molecules such as overlapping molecules, truncated fragments or edge-touching DNA (pixel size 4 nm and scale bar 100 nm). Colors indicate filtration, where molecules touching the edge are black, too short molecules are white, too long molecules are blue and kept molecules are purple. (G) Histogram shows calibration result, colored by filtration; molecules filtered out because they were at an edge (black) too short (white) or too long (blue), as well as the kept lengths (purple) in bp. Distribution is centered around the known length of 1633 bp as a result of the calibration (Methods and Materials). (H) Application of calibration constant to non-filtered DNA segmentations, here shown on 4 nm AFM image 1633 bp plasmids, color bar indicates calculated length of given DNA molecule (scale bars 100 nm), Gaussian fit of the dominant peak found the mean to be 1610 bp with a standard deviation of 143 (N=43).

Ground-truth proximity maps were generated from manual DNA annotations by computing the Euclidean distance to the nearest annotated DNA contour, and subsequently normalizing this distance to [0,1], and then inverting it (1−d) so that backbone pixels (distance 0) have the maximum value and distant background approaches the minimum. Because manual annotation of AFM DNA molecules is time-consuming and therefore yields a limited training set, we substantially expanded the effective dataset through extensive augmentation (Methods and Materials). These augmentations mimic experimental variability in pixel size and signal-to-noise ratio (SNR), allowing the network to generalize to new datasets while reducing the need for additional manual tracings. Beyond generalization across imaging conditions, DNAsight also performs robustly across different AFM instruments without retraining (Figure S3/S4), demonstrated on a previously published dataset^34^.

During inference, DNAsight does not use the predicted proximity maps directly. Instead, they are processed through a multi-step skeletonization pipeline. A threshold is applied to the predicted proximity map to identify candidate DNA regions. Spurious detections are removed by area filtering, and a difference-of-Gaussians (DoG) filter enhances backbone ridges. A combined threshold on the DoG response and the predicted proximity map produces refined binary traces, which are then skeletonized to a one-pixel-wide backbone (Figure 2A/B and S3). These skeletons form the output of S1 and provide continuous centerlines suitable for downstream quantifications.

Segmentation accuracy was evaluated using pixel-distance error metrics (Figure 2C and S1, Methods and Materials), which directly quantify deviations between predicted and manually annotated medial axes. Distances were computed in a directional nearest-neighbor manner, such that the average distance from annotation A to annotation B is not necessarily equal to the reverse comparison. Unlike typical overlap-based scores such as Intersection over Union (IoU)^35^ or Dice^36^, which penalize small lateral offsets and are poorly suited to sparse, one-pixel-wide traces, distance errors directly assess the backbone precision required for downstream analyses.

In Figure 2D, predictions were compared against a manual annotation across a range of SNRs, where each image was assigned a single SNR value computed from the average signal and background statistics of all DNA molecules based on the segmentation mask (Methods and Materials). DNAsight performed robustly under all conditions, with accuracy improving at higher SNRs as expected. Across a range of SNRs from 1.04-2.15, DNAsight achieved 76% of skeleton points within 1.5 pixels of the manual trace on the 10% test dataset (trained on the remaining 90%), indicating that the automated backbone localizes the DNA contour with accuracy comparable to the spatial resolution of the AFM image and the precision of manual annotation.

To place this in context, we compared DNAsight predictions against multiple independent human annotations and annotators (Figure 2E). The resulting inter-annotation distance map is asymmetric due to the directional nature of the metric, which may reflect differences in annotation style. Human annotations can exhibit local variability along the DNA contour, whereas DNAsight produces a comparatively smoother backbone representing regions that are consistently annotated across the training data. As a result, distances from human annotations to DNAsight tend to be smaller, while the reverse comparison can yield slightly larger values because the automated trace does not capture all local deviations present in individual human annotations. Crucially, because independent human annotations do not coincide, zero deviation from any single trace is neither expected nor the appropriate benchmark. Instead, the mean deviation between DNAsight and an individual human annotation falls within the same range as the deviations observed among independent human annotators, indicating agreement at the level of inter-annotator variability.

To further evaluate segmentation accuracy, we benchmarked DNAsight against TopoStats^19^ using three full-field AFM images of DNA-only samples with SNRs of 1.66, 1.75, and 1.81. Each image contained multiple DNA molecules, and segmentation accuracy was similarly quantified as the average pixel deviation between automated traces and manual annotations. DNAsight achieved mean deviations of 0.58, 0.60, and 0.68 pixels, respectively, compared to 2.05, 1.54, and 1.77 pixels for TopoStats (Figure S5). The comparison was limited to the segmentation module of TopoStats, excluding its downstream analysis features. For TopoStats, segmentation parameters were individually optimized for each image to achieve optimal performance, whereas DNAsight operated directly on raw, full-field inputs without parameter adjustment.

To ensure biological interpretability, DNAsight incorporates a DNA length quantification module (Q1). Calibration is first performed on images of DNA molecules of a known length, which are used to calculate a conversion factor from pixels or nanometers to bp. Users can supply either a single image with a known pixel size and DNA length, or multiple images to refine and improve the overall calibration. Considering biological variability, outlier molecules such as truncated fragments, overlapping DNA, or edge-touching molecules can optionally be excluded to improve accuracy (Figure 2F). The length of each skeletonized DNA molecule is then extracted based on Euclidean distance principles, and a calibration constant is calculated from the mean length (Methods and Materials).

This dataset-specific calibration accounts for differences in microscope resolution, scan size, and surface chemistry. When such calibration is unavailable, users can optionally use the canonical B-form DNA value of 0.34 nm/bp^37^ or another value, however internal calibration is recommended as the calibration can vary depending on pixel size and experimental conditions (Figure S6/7). Notably, there is no single optimal pixel size for DNA segmentation or calibration across all experimental systems. Smaller pixel sizes can improve measurement and quantification precision by increasing spatial sampling density, but they typically also reduce the field of view and thereby lower throughput and potentially crop molecules. Conversely, overly large pixel sizes lead to pixelated DNA contours and loss of structural information needed for reliable tracing. Pixel size selection therefore reflects a practical compromise between resolution and sampling depth. The current ML model was trained on images spanning approximately 1–5 nm/pixel, reflecting the practically useful range represented in our data. In practice, a given Q1 calibration is ideally applied to AFM datasets acquired under similar imaging conditions. This is demonstrated in Figure 2G-H, where a calibration derived from 10 images of a 1633 bp plasmid construct (Figure 2G) was applied to an independent image (Figure 2H) acquired under identical conditions but not used in the calibration procedure. As visualized with a colorbar in Figure 2H, both plasmid and linear DNA lengths are accurately determined to be around 1633 bp, where truncated molecules are shorter and overlapping molecules are longer, further validating modules S1 and Q1. Fitting the dominant peak of the resulting length distribution (Figure S8, Methods and Materials) yielded a mean contour length of 1610 bp with a standard deviation of 143 bp (N = 43 molecules), demonstrating that the calibration generalizes well to unseen data. In addition, Q1 flags DNA molecules that touch the image edge, which can be optionally excluded, as they may not represent complete molecules.

Together, segmentation and calibration convert AFM topographs into skeletonized, bp-calibrated DNA backbones at the SM level. This foundation enables robust downstream quantification of chromatin features and could be adapted to other nucleic acid substrates with additional training data.

### Quantification of Supercoiling and Protein-Induced Changes in DNA Spatial Organization

With segmentation established and calibration available when needed, we next moved from backbone tracing to structural insight by applying DNAsight’s spatial organization module (Q2) (Figure 3A). Q2 extracts seven categories of features from skeletonized DNA traces, together providing a comprehensive description of DNA geometry. These include DNA compaction, defined as a rescaled radius of gyration normalized to the DNA length; the number of crossings; tortuosity, defined as the ratio of contour length to end-to-end distance; the frequency of strong bends (default is < 60°^38–40^, but can be adjusted by the user); curvature statistics (mean, standard deviation, minimum and maximum of absolute curvature); persistence length; and degree of elongation (see Methods and Materials for details).

**Figure 3.**
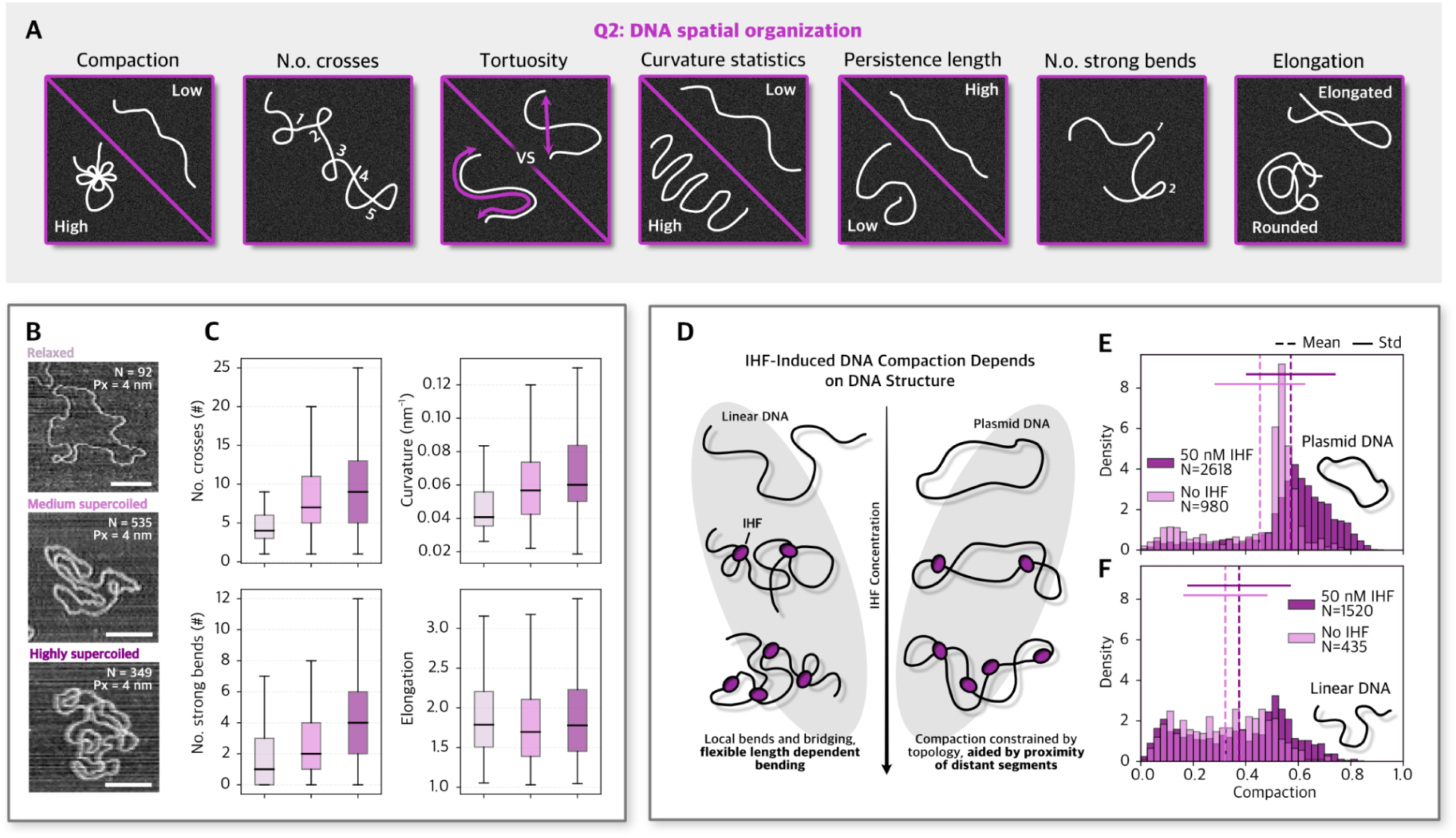
Quantification of DNA spatial organization. (A) Overview of the Q2 module output. From skeletonized DNA traces, DNAsight extracts seven categories of descriptors: length normalized radius of gyration (compaction), number of crossings (self-crossings), tortuosity, curvature statistics, persistence length, frequency of strong bends and a shape descriptor (elongation). (B) Representative images of DNA molecules of a relaxed (light pink), medium supercoiled (pink), and highly supercoiled plasmid (dark pink), pixel size (Px) is 4 nm for all images. (C) Boxplots of number of crossings (relaxed: 5.54 ± 5.25, medium: 8.68 ± 6.71, supercoiled: 10.67 ± 8.00), mean curvature (relaxed: 0.06 ± 0.05, medium: 0.13 ± 0.81, supercoiled: 0.09 ± 0.12), number of strong bends (relaxed: 2.09 ± 2.34, medium: 3.19 ± 3.08, supercoiled: 4.36 ± 3.66) and elongation parameter (relaxed: 1.96 ± 0.65, medium: 1.83 ± 0.60, supercoiled: 1.90 ± 0.61) of relaxed (N=92), medium (N=535), and supercoiled (N=349) DNA. (D) Cartoon of the expected effect of IHF on linear versus plasmid DNA. (E) Quantification of compaction (dashed line, mean; solid line, ±standard deviation), in plasmid DNA with (light pink, mean=0.45±0.17, N=980) and without (dark pink, mean=0.57±0.17, N=2618) IHF. Higher value of compaction indicates stronger compaction. (F) Quantification of compaction in linear (bottom) with (light pink, mean=0.32±0.16, N=435) and without (dark pink, mean=0.38±0.20, N =1520) IHF.

Importantly, Q2 can be applied independently of Q1. Several features, compaction, crossings, tortuosity and elongation, are unitless and directly comparable across datasets. The remaining features, curvature, persistence length and strong bends, depend on spatial scale and require a known pixel size to be comparable across images of varying pixel size. Calibration (Q1) is only needed when relating geometric descriptors to DNA length in bps, where it adds interpretive value by connecting nanometer-scale measurements to sequence length. Thus, Q2 functions as a flexible module: it can be run entirely on its own for relative comparisons, requires pixel size for cross-dataset comparability, and can leverage calibration when length-dependent interpretation is desired.

To demonstrate the breadth of Q2, we highlight two representative examples. DNA topology can vary through torsional strain that twists the double helix upon itself, producing structures known as supercoils^41,42^. Negative supercoiling generally promotes local DNA unwinding and looping, whereas relaxed molecules adopt more extended conformations. We applied DNAsight to DNA plasmids classified with varying levels of supercoiling (as shown in a previous publication^43^), namely as relaxed, medium supercoiled, or highly supercoiled (Figure 3B), imaged with a pixel size ranging from 2-4 nm. Here, we show four of the aforementioned Q2 categories. We find that the number of crossings, strong bends, and curvature expectedly increase with supercoiling, whereas elongation remains largely unchanged (Figure 3C, representative image outputs in Figure S9).

Second, we applied Q2 to study protein-induced compaction of DNA using IHF. IHF is a small, heterodimeric DNA-binding protein from *Escherichia coli* that introduces sharp bends in DNA and plays a crucial role in organizing bacteria chromatin architecture and regulating diverse DNA transactions such as replication, recombination, and transcription^41^. This process is expected to differ between linear and plasmid topologies because circular DNA is topologically constrained, whereas linear DNA is not (Figure 3D). Here, we applied the compaction feature which is designed to range between 0 (low compaction) and 1 (high compaction) to images of pixel sizes ranging from 3-4 nm. Before the addition of IHF, the linear DNA showed a more varied compaction, where the plasmid compaction displayed less variation with a concentrated peak around 0.55 (Figure 3E). Upon addition of IHF, plasmids shifted markedly toward more compact configurations (mean compaction 0.45±0.17 to 0.57±0.17; Δ_mean_ = +0.12, 26%), while linear DNA exhibited only a modest increase (mean compaction 0.32±0.16 to 0.38±0.20; Δ_mean_ = +0.06, 13%) (see representative image outputs in Figure S10). Although the distributions remained broad (standard deviation shown), both plasmid and linear DNA showed significant IHF-induced increases in compaction (two-sided Mann–Whitney U test, plasmid: p=7.5×10^-110^; linear: p=5.5×10^-8^). Whereas the increase was larger for plasmids than for linear DNA (difference in mean shifts = 0.065; permutation test, p=1.0×10^-4^). This can be rationalized by DNA topology and length constraints. Plasmids provide a closed circular template that keeps distant segments in closer proximity, so IHF-induced bends and bridging can more readily stabilize compact configurations. By contrast, for linear DNA, particularly at the 1.7 kb length used here, distal segment encounters are expected to be less frequent, consistent with reports that bridging efficiency tends to increase with contour length in the kilobase regime^44–46^.

Together, these examples illustrate how Q2 can resolve both global changes in DNA compaction and local structural perturbations such as bending and curvature. By combining seven complementary descriptors rather than relying on a single metric, DNAsight captures the complexity of chromatin folding states and facilitates quantitative comparisons across datasets and conditions.

### Automated Detection of DNA Loop-Like Structures Reveals Condition-Dependent Enrichment of CTCF-Proximal Loops

Moving beyond the aforementioned geometric features, many biologically relevant rearrangements arise locally through DNA looping and bridging, for example when SMC complexes or transcription factors bring distant DNA segments, and their regulatory sites, into proximity^47–49^. To quantify such events, DNAsight includes a dedicated loop-quantification module (Q3), which detects loop-like closed structures along DNA molecules (Figure 4A). Computationally, Q3 operates on the skeletonized DNA traces produced in S1 and identifies regions where non-adjacent segments of the backbone approach each other within a defined spatial threshold and form a closed path along the skeleton. Loop-like structures are operationally defined as closed paths in the skeleton graph, corresponding to backbone segments that connect to form a closed contour. Each detected loop-like structure is reported with its contour length and position relative to the closest DNA end, enabling systematic quantification.

**Figure 4.**
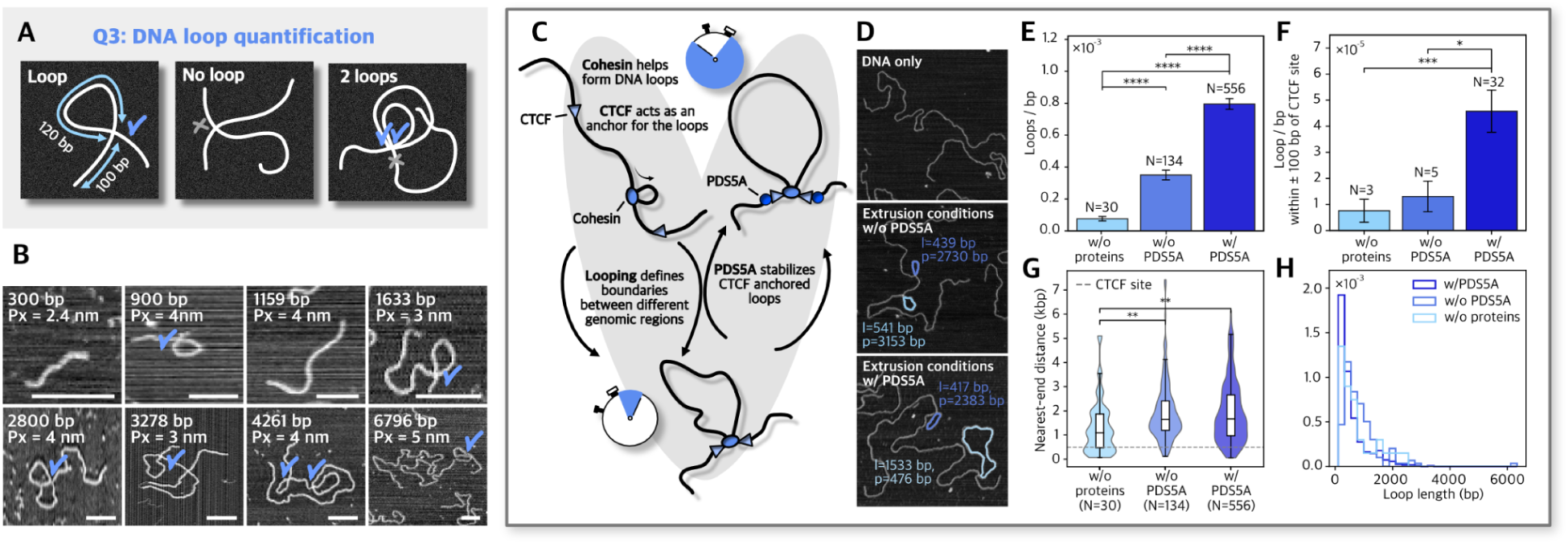
Detection and quantification of loop-like structures in DNA and cohesin–CTCF reactions. (A) Overview of logic of the Q3 module. Skeletonized DNA traces are analyzed to detect closed contours, with each loop reported by position and length. Loop-like structures are distinguished from simple overlaps when the contour is open. (B) Representative AFM images of DNA molecules of varying length, all in the absence of protein, blue tick marks indicate loops detected by Q3, pixel size is given as Px. (C) Cartoon illustrating the working model motivating analysis of loop-like structures in cohesin-CTCF reactions with and without PDS5A. (D) Representative raw image of DNA molecule of each experimental condition, DNA only, extrusion conditions w/o PDS5A and extrusion conditions w/ PDS5A, showing Q4 output of loop outlines with labelled loop length (l) and position (p), pixel size is 3 nm for all images. (E) Loop frequency per bp for naked DNA with no proteins present (N=30), then for CTCF, cohesin-STAG1, NIPBL, and ATP in the absence (N=134) or presence (N=556) of PDS5A. (F) Loop frequency per bp with +/-100 bp of the CTCF binding site for naked DNA (N=3) and with proteins with (N=5) and without PDS5A (N=32). (G) Violin and boxplots of the nearest-end distance or position of loop-like-structures on DNA for naked DNA and with proteins with and without PDS5A. (H) Histograms show distribution of loop lengths in bp for naked DNA and with protein with and without PDS5A. Significance stars were assigned from two-sided Mann–Whitney U tests with Bonferroni correction for multiple comparisons. Each p-value was multiplied by the number of tested pairs, and stars were displayed according to adjusted significance thresholds: *(p < 0.05), **(p < 0.01), ***(p < 0.001), and ****(p < 0.0001); comparisons not significant after correction were left unmarked.

It is important to distinguish these loop-like structures from other forms of apparent self-contact. Crossings, reported in Q2, correspond to local crossings of the DNA backbone that do not necessarily form a closed path. Loop-like structures, in contrast, are only annotated when the skeleton topology contains a complete closed path. This algorithmic distinction provides a consistent and reproducible way to separate closed loop-like configurations from simple crossings (Figure 4A). As illustrated in Figure 4B, loop-like structures can also arise spontaneously in the absence of bound proteins, particularly in longer DNA molecules that have more opportunities for self-contact. Thus, not all closed contours necessarily reflect protein-mediated looping. For this reason, loop quantification (Q3) should be interpreted relative to appropriate controls unless anchoring proteins can be directly visualized at the loop base. Q3 also records the mean and standard deviation of the intensity of each DNA contour and loop anchoring point, which can support interpretation in datasets where anchoring proteins are detectable. In practice, however, anchoring proteins are not always reliably visible in AFM images because of imaging conditions or because they become displaced or dissociate during sample preparation or imaging, particularly in the absence of cross-linking or other stabilizing treatments. For the datasets presented here, we therefore focus on comparison to appropriate no-protein controls as the basis for interpreting loop-like structures.

To move from general loop-like geometries toward a defined biological system, we reconstituted DNA substrates under conditions that mimic cohesin-mediated loop extrusion. Previous studies have suggested that PDS5 stabilizes cohesin-CTCF-DNA loops by reinforcing cohesin residence and anchoring at CTCF sites^27,50^. To examine this proposed stabilization through direct AFM visualization, we designed a symmetric 8 kbp DNA construct containing 2 CTCF binding sites 493 bp from each end and performed in vitro looping reactions using an established minimal system for cohesin loading^51,52^, extrusion, and stalling: CTCF, cohesin-STAG1-NIPBL (hereon referred to as “cohesin”), and ATP, with or without subsequent addition of PDS5A (Figure 4C). To establish a baseline for spontaneous DNA looping, we also conducted a DNA-only control under the same conditions. All conditions were imaged with a 3 nm pixel size.

Across these conditions (Figure 4D), DNAsight detected a progressive increase in the frequency of loop-like structures occurring anywhere along the DNA molecules as proteins were added. The inclusion of CTCF and cohesin led to a 4.6-fold increase in detected loop-like structures relative to the DNA-only baseline, with a further 2.3-fold increase upon addition of PDS5A (Figure 4E). The increased frequency of loop-like structures in the presence of PDS5A is consistent with, but does not by itself establish, a contribution of PDS5A to loop stabilization under these reconstitution conditions^27^. A similar trend was observed for loop-like structures positioned within a ± 100 bp window around expected CTCF-sites. Because loop position is defined relative to the nearest DNA end, this enrichment should be interpreted cautiously, as nonspecific loops can also fall within this positional window by chance. The frequency of CTCF-site-proximal loop-like-structures increased 1.7-fold from the DNA-only baseline to the cohesin-CTCF condition, and a further 3.5-fold increase upon the addition of PDS5A (Figure 4F). Notably, CTCF-site-proximal loop-like-structures constituted only a small fraction of the total (Figure 4E-F). Even when CTCF, cohesin, and PDS5A were present, more than 90% of loop-like-structures occurred away from the CTCF site, consistent with a substantial population of non-CTCF-anchored loop-like-structures. DNA molecules overlapping the image edge, shorter than 5 kbp or longer than 10 kbp were excluded to remove partially imaged, truncated or overlapping DNA molecules, and loop counts were normalized to total detected DNA length to account for differences in image area and DNA density. To reduce duplicate counting of nested or partially overlapping closed contours on the same DNA (exemplified in Figure 4A right), highly overlapping loop candidates were filtered as described in Methods; the corresponding analysis without this filter is shown in Figure S11. These results show that, under the reconstitution conditions used here, PDS5A-containing samples exhibit a higher frequency of detected loop-like structures, increasingly so within a CTCF-proximal window (Figure 4G-F; see additional representative images with overlaid loop detection in Figure S12). However, because anchoring proteins were not directly visualized and most detected loop-like-structures were not CTCF-proximal, these observations should be interpreted as suggestive rather than definitive evidence for specific cohesin-CTCF loop stabilization. Lengths of loop-like structures ranged from several hundred to a few thousand base pairs (Figure 4H), consistent with a heterogeneous population of DNA self-contacts and protein-associated loop-like configurations. Using an unbinned kernel density estimate, the most frequent loop lengths were 440.1 bp (95% bootstrap CI: 357.5-549.0 bp) for DNA only, 569.7 bp (95% bootstrap CI: 474.4-657.9 bp) for CTCF-cohesin, and 274.1 bp (95% bootstrap CI: 255.2-294.9 bp) for CTCF-cohesin-PDS5A. If loops were anchored between the two CTCF sites on the symmetric construct, they would be expected to span approximately 7 kbp. The substantially shorter most frequent loop lengths therefore suggest that the detected loop-like structures predominantly reflect a heterogeneous mixture of partial looped states and nonspecific overlaps rather than a dominant population of fully site-to-site anchored loops.

Together, these experiments demonstrate how Q3 provides systematic, quantitative measurements of the number, size, and position of loop-like structures. Nevertheless, appropriate control samples are essential for interpreting these results, as they establish the baseline frequency of loop-like structures and spatial organization of DNA in the absence of protein-mediated effects.

### Quantifying Protein-DNA Assemblies and Nucleosome Architecture

In addition to DNA segmentation (S1), DNAsight includes a protein segmentation module (S2) for detecting protein features across a broad range of sizes and morphologies, from compact nucleosome-like particles to large multi-protein assemblies (Figure 5A). To accurately segment protein assemblies that span a wide range of sizes and intensities, DNAsight employs two segmentation approaches. First, the large-cluster segmentation is designed for broad, continuous protein assemblies of various shapes. It applies a smoothed intensity-based segmentation, followed by region growing and area filtering, to delineate spatially contiguous, high-intensity regions while excluding small speckles and noise. In contrast, the small-spot segmentation targets compact foci like nucleosomes. It utilizes principles from single-particle detection with local thresholding around each detection to generate a refined mask for every spot, followed by geometric filtering based on area, circularity, and eccentricity. Together, these two strategies allow S2 to sensitively detect both sparse puncta and extended assemblies while providing a common set of descriptors, including position, area, and intensity, for downstream analysis (Methods and Materials).

**Figure 5.**
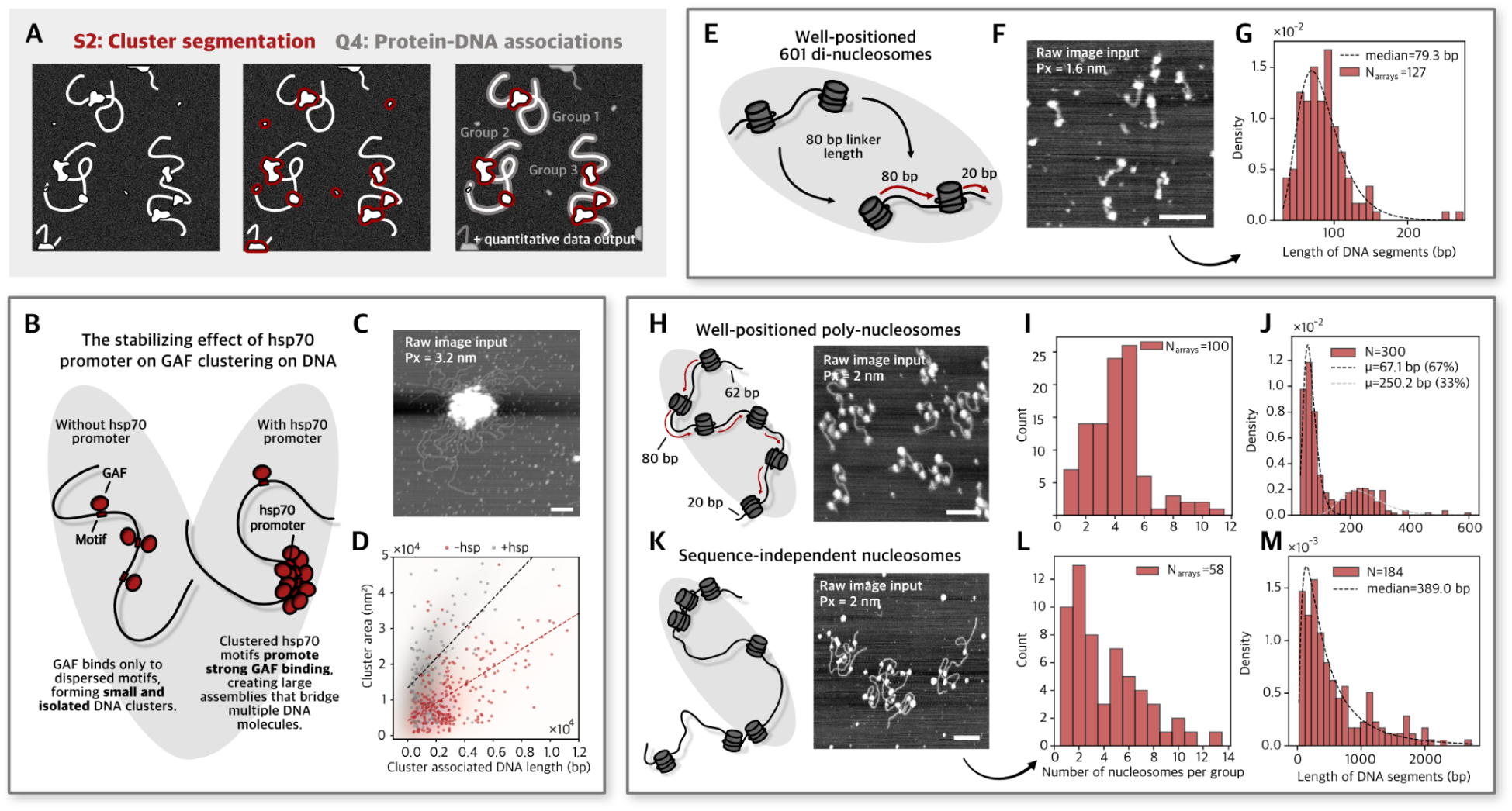
Quantification of protein–DNA associations and nucleosome spacing. (A) Overview of Q4. DNA (S1) and protein (S2) segmentation masks are integrated to quantify protein–DNA associations, filtering can optionally be applied to remove edge and/or too small clusters. (B) Cartoon demonstrating the hypothetical effect of the *hsp70* promoter on GAF clustering, with more clustering happening in the presence of the *hsp70* promoter. (C) Representative AFM images showing GAF binding to DNA with *hsp70* motifs, producing a large protein cluster (pixel size 3.2 nm). (D) Quantification of cluster area versus length of associated DNA with (N=130) and without (N=170) the presence of the *hsp70* promoter. Groups at the edge, too small clusters as well as groups containing more than one cluster were filtered out, see details in Methods and Materials. (E) Cartoon of di-nucleosome construct with 80 bp linker length. (F) Representative AFM image of di-nucleosomes (pixel size 1.6 nm). (G) Distribution of of nucleosome-associated DNA segment lengths fit with a lognormal model, yielding a median spacing of 79.3 bp using a lognormal fit with 95% bootstrap CI: 74.5-84.7 bp (N=161), for groups containing two nucleosomes. (H) Left: cartoon of the hexa-nucleosome construct with 80 bp linker length, with 62 and 20 bp at each end. Right: Representative AFM image of hexa-nucleosomes (pixel size 2 nm). (I) Distribution of number of nucleosomes per group for hexa-nucleosome constructs (N=100). (J) Distribution of nucleosome-associated DNA segment lengths in bp for hexanucleosome construct. Distribution is fit with a double lognormal mixture model (supported by BIC, Methods and Materials), with means of 67.1±23.2 bp and 250.2±76.4 bp, representing 67% and 33% of the displayed data, respectively (N=258). (K) Left: cartoon of sequence-independent nucleosome construct (DNA construct lacking 601 positioning sequences) with unknown number of nucleosomes and linker length. Right: Representative AFM image of sequence-independent nucleosomes (pixel size 2 nm). (L) Distribution of the number of nucleosomes of sequence-independent nucleosome constructs (N=58). (M) Distribution of nucleosome-associated DNA segment lengths in bp for sequence-independent nucleosome constructs (N=184), fit with a single lognormal fit with median length of 389.0 bp and 95% bootstrap CI: 323.76-482.90 bp.

The protein-DNA association quantification module (Q4) then integrates the DNA masks from S1 with the protein masks from S2 to quantify protein-DNA associations, including colocalization, binding position along DNA, spacing between protein features, and cluster statistics relative to DNA length or position. As a demonstration, we applied this S2-Q4 workflow to analyze GAF binding on linearized DNA substrates with and without the *Drosophila hsp70* promoter, which contains several GAF-binding sites (Figure 5B), here using the large-cluster segmentation mode. The complexes were imaged with pixel sizes ranging from 2-4 nm. GAF is a sequence-specific transcription factor that recognizes GA-rich motifs and promotes the formation of accessible chromatin domains at heat-shock promoters such as *hsp70*^52,53^. Its ability to multimerize and recruit additional factors is expected to drive cooperative clustering on DNA containing multiple GAF-binding sites^54^. To ensure reliable linking between clusters and DNA, we restricted the analysis to connected DNA-cluster groups, defined as DNA and protein-cluster mask components that spatially overlap after applying a 3-pixel dilation to the cluster mask. For analysis of the GAF constructs, we further restricted the dataset to groups containing a single protein cluster. This was done to ensure unambiguous matching of DNA to a single GAF cluster when quantifying the relationship between cluster size and recruited DNA. In the absence of the clustered motifs on the *hsp70* promoter, GAF bound sparsely, producing smaller, compact protein clusters (representative image outputs in Figure S13). By contrast, DNA carrying the clustered sites on *hsp70* promoter displayed large GAF assemblies bridging multiple DNA molecules (Figure 5C). Quantitative analysis confirmed that the presence of *hsp70* promoter shifted the distribution toward larger cluster areas (Figure 5D), increasing the mean cluster area 1.9 times (Figure S14), consistent with cooperative GAF recruitment to promoter-associated DNA elements. While cluster size increased substantially in the presence of the *hsp70* promoter, the length of detectable DNA, estimated in number of bp, was 1.4-fold higher in constructs lacking the *hsp70* promoter (Figure S14). This difference likely reflects increased DNA compaction within large GAF clusters formed on *hsp70*-containing substrates, which reduces the fraction of DNA that remains visible and quantifiable. Nevertheless, in both conditions we observed a positive relationship between cluster size and the amount of recruited DNA (Figure 5D).

Q4 can also be used to quantify nucleosome positioning from AFM images. Nucleosomes, the fundamental repeating units of chromatin, consist of DNA wrapped around a histone octamer, connected by short linker DNA segments. Here, we used the cluster segmentation for smaller clusters (see representative segmentation in Figure S15) and analyzed well-positioned di-nucleosome (394 bp) and poly-nucleosome (1364 bp) constructs. To resolve these compact constructs, we imaged them at comparatively small pixel sizes ranging from 0.6-2 nm. These were assembled using the 601 positioning sequence, with the poly-nucleosome containing six 601 sites separated by defined 80 bp linker lengths. For di-nucleosome constructs (Figure 5E-F) containing two detected nucleosomes (see unfiltered data in Figure S16), with a median of 79.3 bp from lognormal fitting (95% bootstrap CI: 74.5-84.7 bp; Figure 5G). Although the distribution of individual measurements was broad (standard deviation = 35.0 bp), the central tendency remained close to the expected linker length. The segment length is calculated as the distance of the segmented DNA between the outer edges of segmented nucleosomes (i.e., an edge-to-edge distance), as only exposed DNA is visible. We here characterize the full constructs, including visible terminal segments at the construct ends, although such end-associated segments can be excluded from the output by filtering if desired (Figure S17). Notably, the segmented nucleosome size varied across particles, prompting us to test whether the lengths of nucleosome-associated DNA segments was associated with mean nucleosome size. For the di-nucleosome constructs, we observed no significant Spearman correlation between these variables (Figure S18), indicating that variation in segmented nucleosome size is unlikely to be a major contributor to the measured linker-length distribution.

Poly-nucleosome assemblies (Figure 5H) were analyzed after excluding incomplete and overlapping DNA molecules to focus on representative, interpretable groups. Specifically, DNA groups touching the image boundary were removed, as these correspond to partial molecules. In addition, groups with total detected DNA lengths shorter than two linker lengths (160 bp) were excluded to eliminate severely truncated constructs. To minimize contributions from overlapping assemblies, we further restricted the analysis to groups with total detected DNA lengths not exceeding 629 bp, corresponding to the expected exposed DNA (482 bp) plus one fully unwrapped nucleosome (147 bp). This upper bound was chosen to retain assemblies missing a single nucleosome, which were prevalent in the data, while excluding larger aggregates arising from molecular overlap. After applying these filters, the poly-nucleosome constructs exhibited a mean of 4.2±2.0 nucleosomes per group, with the most frequent value being 5 (Figure 5I, see unfiltered distributions in Figure S16). A small number of groups contained more than the theoretical maximum of six nucleosomes, which likely arise from residual overlapping molecules or local noise and imaging artifacts that were not fully removed by the filtering criteria. Here, the distribution of nucleosome-associated DNA segment lengths was broad and bimodal (supported by Bayesian Information Criterion (BIC); Methods and Materials), with mean values of 67.1 bp and 250.2 bp (Figure 5J). Notably, the two fitted components differed substantially in their spread, with an approximate standard deviation of 23.2 bp for the shorter component and 76.4 bp for the longer component. This indicates that the longer-linker population spans a wider range of measured segment lengths, while the shorter-segment population is more concentrated around its fitted mean. The mean of the shorter component fell below the nominal 80 bp linker length, which may reflect that the construct contains 20 and 62 bp terminal flanking segments, which can shift apparent linker distances toward shorter values. In addition, the hexa-nucleosome construct contains multiple nucleosomes on a longer DNA molecule and is therefore more prone than the di-nucleosome construct to local overlap or partial obscuration of the DNA by neighboring nucleosomes. This may further contribute to underestimation of visible linker lengths. However, the fitted spread still places this component within range of the expected 80 bp spacing. The broad spread of the longer component is consistent with a heterogeneous ensemble of conformations, likely including different degrees of unwrapping, displacement, or absence of individual nucleosomes within the construct, rather than a single discrete structural state. One possible contribution to this population is loss or substantial displacement of an endmost nucleosome, particularly at the 20 bp flank, which would give an expected exposed distance of approximately 247 bp and is therefore close to the fitted mean of 250.2 bp.

We similarly applied Q4 to a DNA construct lacking 601 positioning sequences (hereon referred to as “sequence-independent nucleosomes”), using a 4361 bp template (Figure 5K), imaged with pixel sizes ranging from 1.6-2 nm. The number of nucleosomes per group was more heterogeneous than for the hexa-nucleosome construct, with a median of 3 and mean of 4.17±2.91 nucleosomes and a long-tailed distribution (Figure 5L). The distribution of nucleosome-associated DNA segment lengths was shifted toward substantially longer values relative to both di-and poly-nucleosome constructs, with a lognormal fit yielding a median length of 389.0 bp (95% bootstrap CI: 323.76-482.90 bp, Figure 5M).

Together, these analyses demonstrate that DNAsight can extract structural parameters of chromatin fibers, including nucleosome repeat lengths and heterogeneity, without requiring labeling or external markers.

## DISCUSSION

This work establishes DNAsight as a unified framework for quantitative analysis of AFM images of DNA and chromatin. By combining ML-based segmentation with modular quantification, DNAsight converts nanoscale AFM topographs into biologically meaningful parameters such as DNA length, spatial organization, looping, nucleosome spacing, and protein clustering. The framework operates robustly across diverse imaging conditions, including variations in pixel size, SNR, DNA length, and substrate type, enabling analysis of both linear and plasmid DNA without per-image parameter tuning. Benchmarking against TopoStats confirmed improved segmentation accuracy and human-level agreement with manual backbone tracing, validating DNAsight as a scalable solution for systematic analysis of chromatin organization. Because the segmentation benchmark relies on manual annotations, the reported agreement is inherently relative to expert human tracing rather than an absolute physical ground-truth benchmark. Nevertheless, training on many annotated images allows DNAsight to capture the contour features most consistently identified by human annotators while reducing the influence of local irregularities in individual traces.

A broader consideration is the quantitative precision achievable with automated AFM analysis. Although DNAsight provides standardized and reproducible structural readouts, the effective resolution depends not only on the analysis itself but also on the quality and information content of the underlying AFM data, including pixel size, signal-to-noise, tip convolution, calibration quality, surface effects, and molecular heterogeneity. Thus, the current limits on precision are likely driven more by experimental acquisition than by the analytical framework itself. In its current form, DNAsight is well suited to detecting robust population-level differences in features such as nucleosome spacing or loop size, whereas interpretation of very small differences, such as base-pair-scale shifts in nucleosome spacing^55^, should be made with appropriate caution. Larger sample sizes can improve the precision of population estimates, but not the underlying per-molecule measurement uncertainty. Further gains in resolution are therefore likely to come from improved imaging, although continued analytical advances may also further improve precision.

Beyond segmentation, DNAsight introduces analytical capabilities that were not previously readily accessible for AFM data. Its modular design integrates structural and biochemical perspectives within a single pipeline, enabling quantification of local curvature, loop-like structures, and protein-DNA associations in the same dataset. Applying DNAsight to reconstituted chromatin systems revealed topology-dependent DNA compaction by IHF, condition-dependent changes in loop-like DNA structures in cohesin-CTCF-PDS5A reactions, and *hsp70* promoter-dependent clustering of GAF on DNA. Likewise, nucleosome-spacing analysis extracted precise linker-length distributions directly from raw AFM images, demonstrating how DNAsight enables biologically interpretable measurements that were previously limited to small-scale or manual analyses.

This interpretability should, however, be considered in light of how the readouts are derived. DNAsight quantifies topology from 2D backbone-proximity maps and skeletonized segmentations, enabling robust and scalable analysis across heterogeneous AFM datasets, but without fully exploiting the height information in the underlying topographs. Consequently, some features are represented as reduced topological descriptors rather than complete structural reconstructions: (1) crossings in highly supercoiled DNA may underestimate apparent contour length, (2) loop-like structures may sometimes reflect overlap rather than protein-mediated anchoring, and (3) DNA embedded within dense protein clusters may be underrepresented in length-based measurements.

Furthermore, certain analysis choices are pragmatic by design. For example, loop position is defined relative to the nearest DNA end, which is useful for comparing molecules but does not preserve directionality, limiting assignment of individual loops to specific genomic orientations or binding sites. These constraints are especially relevant for mechanistic interpretation of single events. Thus, in the cohesin-CTCF-PDS5A experiments, the enrichment of CTCF-proximal loop-like structures is best interpreted as suggestive rather than definitive. Overall, these considerations underscore both the analytical utility of DNAsight and the importance of interpreting AFM-derived topological features in the context of the structural information available in the images and, where possible, complementary biochemical controls.

By unifying segmentation, calibration, and quantification within a user-friendly graphical interface, DNAsight enables reproducible, high-throughput AFM analysis of chromatin across experimental systems. This integration bridges molecular imaging with quantitative structural biology, allowing large-scale comparative studies and uncovering organizational features of DNA and DNA-protein assemblies that were previously inaccessible, even in existing AFM datasets.

## ACKNOWLEDGMENTS

We thank members of the Hatzakis and Ha laboratories for helpful discussions and feedback, and we acknowledge the support of shared facilities and staff at the authors’ institutions. During the preparation of this work the authors used OpenAI’s ChatGPT for assisting with programming and refining writing. After using this tool, the authors reviewed and edited the content as needed and take full responsibility for the content of the published article. This article is subject to HHMI’s Immediate Access to Research policy, which requires that this article be made publicly available as initial and revised preprints deposited on a designated preprint server under a CC BY 4.0 license.

## AUTHOR CONTRIBUTIONS

E.W.S., S.P., J.B.K., and T.H. conceived the project. E.W.S. designed and developed the DNAsight analysis framework, including the core codebase, ML-based segmentation, and downstream quantification modules, with contributions and guidance from J.B.K. S.P. and T.-W.L. performed the AFM experiments and imaging, and led experimental data collection; T.-W.L. led initial development/optimization of AFM surface-preparation protocols. P.J.M. and S.R. designed and prepared samples as well as assisted with the imaging for the looping experiments. R.U.-M. designed nucleosomal DNA constructs, prepared nucleosomes, and developed the nucleosome reconstitution protocol. F.R. expressed and purified IHF, and J.H. generated DNA supercoiling constructs. I.F.D. expressed, purified and labelled human CTCF and mutants. L.C. expressed and purified cohesin and PDS5A, and M.O.-V. expressed and purified NIPBL. M.Y. and X.A.F. expressed and purified GAF and the related constructs. Manual annotations for ML training and performance evaluation were produced by E.W.S., S.P., P.J.M., R.U.-M., and S.G.. E.W.S. performed quantitative analysis and statistical evaluation. E.W.S. S.P. R.U.-M. P.J.M, S.R. and T.H. contributed to interpretation of the results and biological conclusions. TopoStats benchmarking was carried out by E.W.S., with J.Z. providing expertise and technical support that enabled implementation and interpretation. N.S.H. contributed advice and ongoing support throughout the project. L.F., S.M.V., J.-M.P., J.B. and C.W. provided expertise and support for the collaboration. The work was supervised by J.B.K. and T.H.. E.W.S. wrote and created visuals for the manuscript with input from S.P., R.U.-M., P.J.M., S.R., J.B.K., and T.H.

## DATA AVAILABILITY STATEMENT

All data underlying this study are publicly available in Zenodo under record 18953785. The frozen software release associated with this publication is available in Zenodo under record 1895407. The actively maintained development version of the DNAsight source code is available on GitHub at kirkegaardlab/dnasight.

