## Supporting Information for "A Modular Framework for Automated Segmentation and Analysis of AFM Imaging of Chromatin Organization"

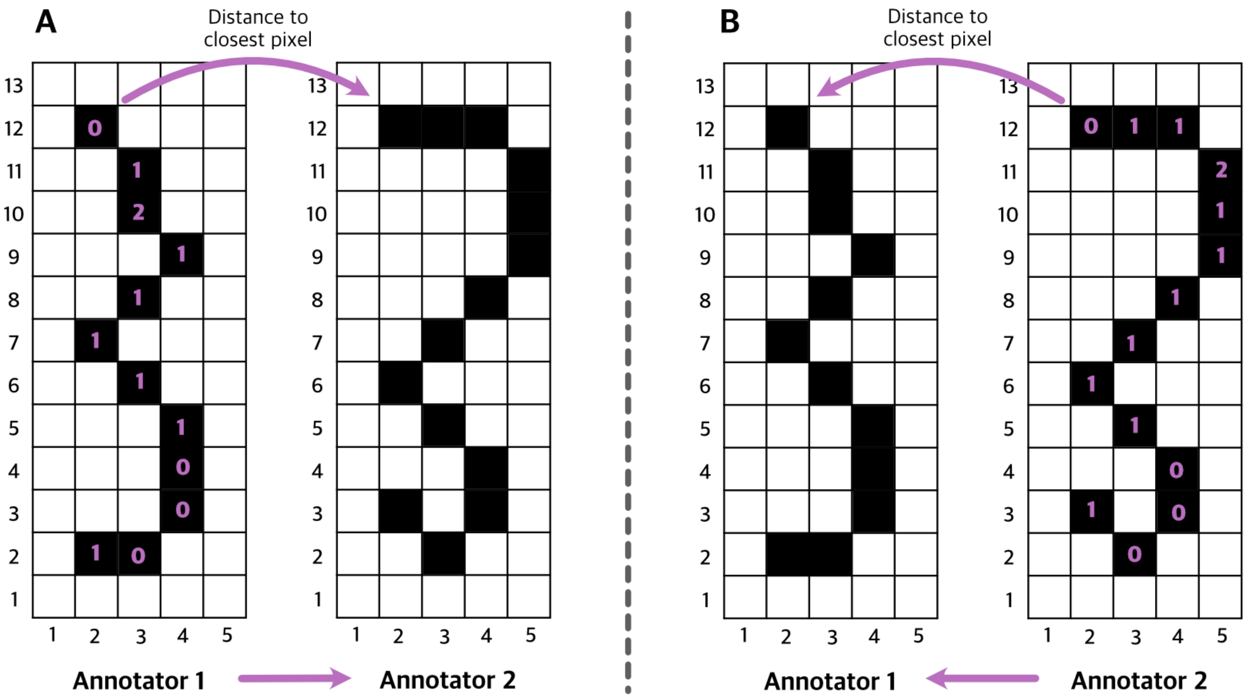


**Figure S1. DNA segmentation performance metric for average pixel variation between annotators.** (A) Cartoon displaying the distance to the closest pixel from annotator 1 to annotator 2. (B) Same depiction as in (A), but the other direction from annotator 2 to annotator 1.

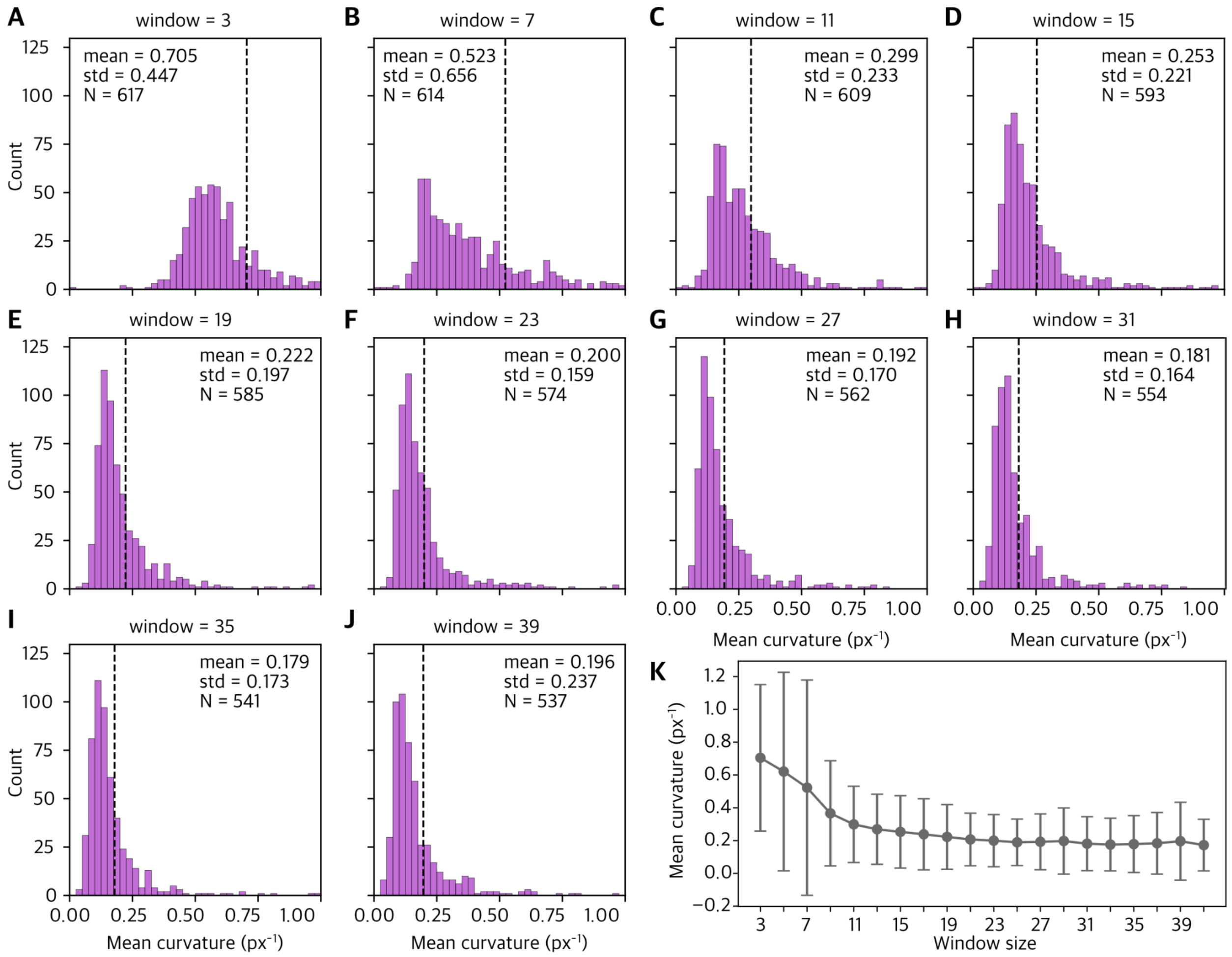


**Figure S2. Sensitivity of curvature estimates to Savitzky–Golay smoothing window.** (A-J) Histograms show the distribution of mean curvature values of medium supercoiled DNA molecules using different Savitzky–Golay smoothing-window sizes (3-39, window sizes indicated in each panel). Dashed vertical lines mark the mean curvature for each setting. N denotes the number of valid molecules retained for the given smoothing window size setting. (K) Mean and standard deviation of the overall mean curvature as a function of Savitzky–Golay smoothing-window size.

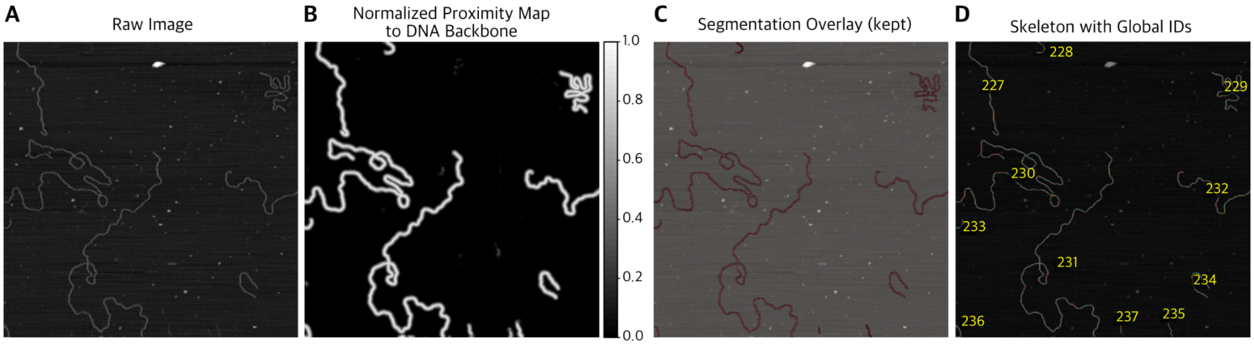


**Figure S3. Automated segmentation and skeletonization workflow for unannotated AFM images (RIBM SS-NEX Ando HS-AFM).** (A) Raw image input (pixel size 3 nm). (B) Predicted normalized backbone-proximity map output by the trained U-Net, where values range from 0 to 1 and brighter pixels indicate higher proximity to the DNA centerline (maximum on the backbone, decreasing with distance). (C) Watershed-segmented regions from (B) that pass the area filter are displayed in semi-transparent red on the raw image, representing DNA molecules retained for further analysis. (D) Final skeletons extracted using DoG filtering and thresholding, restricted to the kept regions. Each distinct DNA contour is assigned a unique global identifier (colored overlay, yellow numbers), ensuring non-overlapping IDs across all images in the dataset.

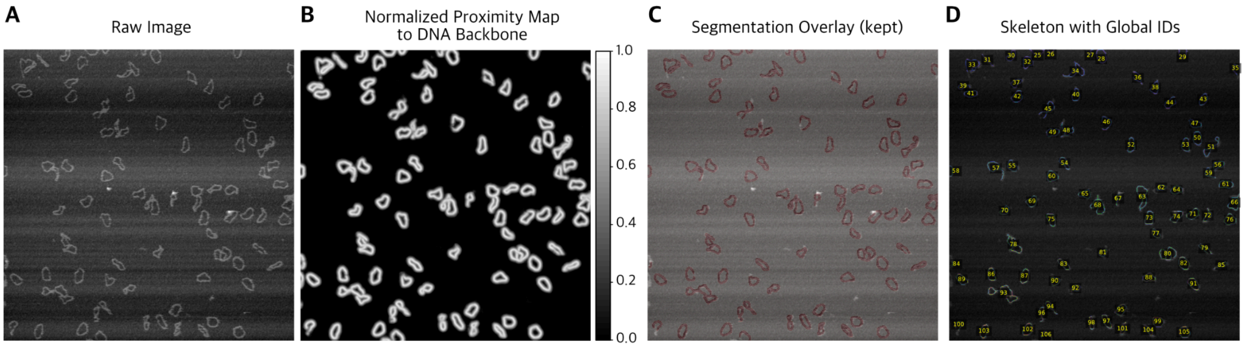


**Figure S4. Automated segmentation and skeletonization applied to commercial Bruker HS-AFM data.** (A) Raw image input (pixel size ≥1 nm). (B) Predicted normalized backbone-proximity map output by the trained U-Net, where values range from 0 to 1 and brighter pixels indicate higher proximity to the DNA centerline (maximum on the backbone, decreasing with distance). (C) Watershed-segmented regions from (B) that pass the area filter are displayed in semi-transparent red on the raw image, representing DNA molecules retained for further analysis. (D) Final skeletons extracted using DoG filtering and thresholding, restricted to the kept regions. Each distinct DNA contour is assigned a unique global identifier (colored overlay, yellow numbers), ensuring non-overlapping IDs across all images in the dataset. Data used from previous publication[1].

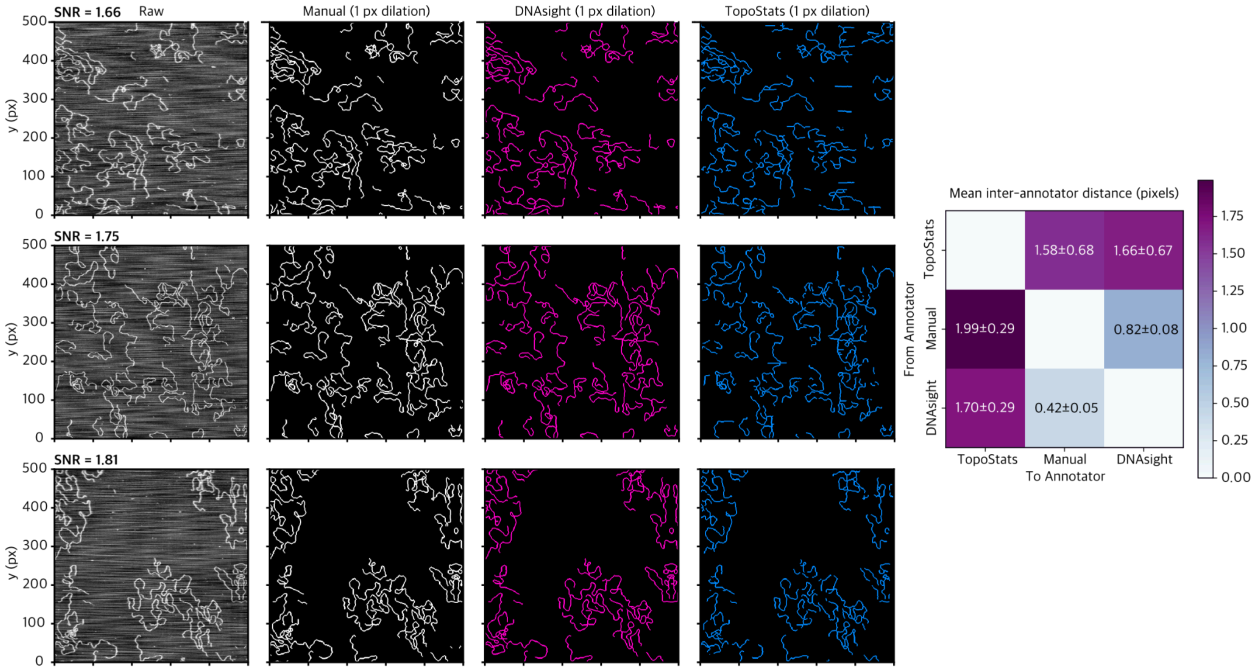


**Figure S5. Benchmarking DNAsight against TopoStats.** First column, 3 raw images of only DNA with pixel size 4 nm and SNR 1.66, 1.75 and 1.81. Second column, manual DNA segmentation. Third column, DNA segmentation using DNAsight. Fourth column, DNA segmentation using TopoStats. Matrix shows inter-pixel-distances between manual, DNAsight and TopoStats segmentations.

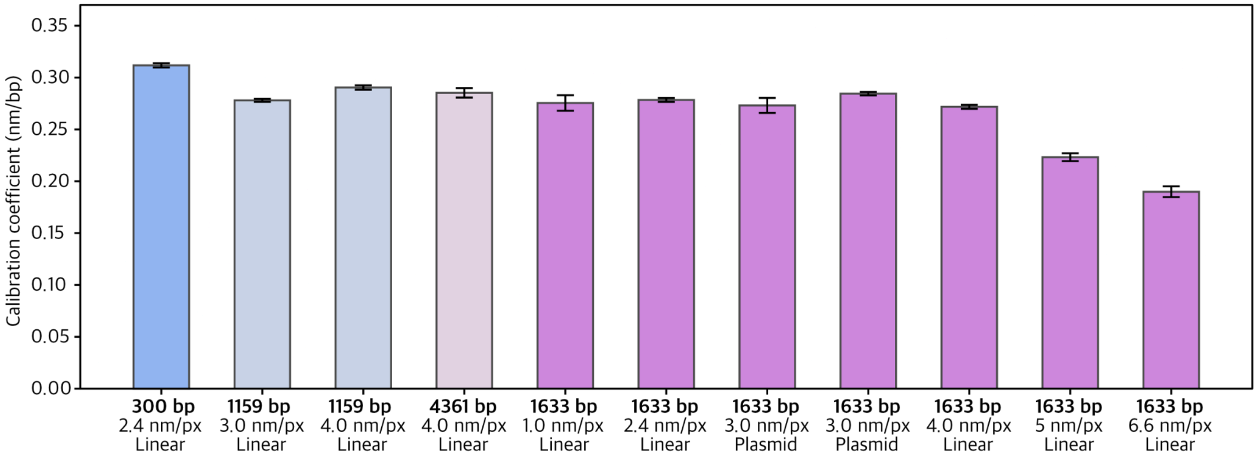


**Figure S6. Calibration coefficient determined by DNAsight module Q1 for different pixel sizes and DNA constructs.** From the left: 300 bp linear construct measured with pixel size of 2.4 nm, 1159 bp linear construct measured with pixel size of 3 nm and 4 nm, 4361 bp linear construct measured with pixel size of 4 nm, 1633 bp linear construct measured with pixel size of 1, 2.4 and 3 nm, 1633 bp plasmid construct measured with pixel size of 3 nm and 4 nm and 1633 bp linear construct measured with pixel size of 5 nm and 6.6 nm.

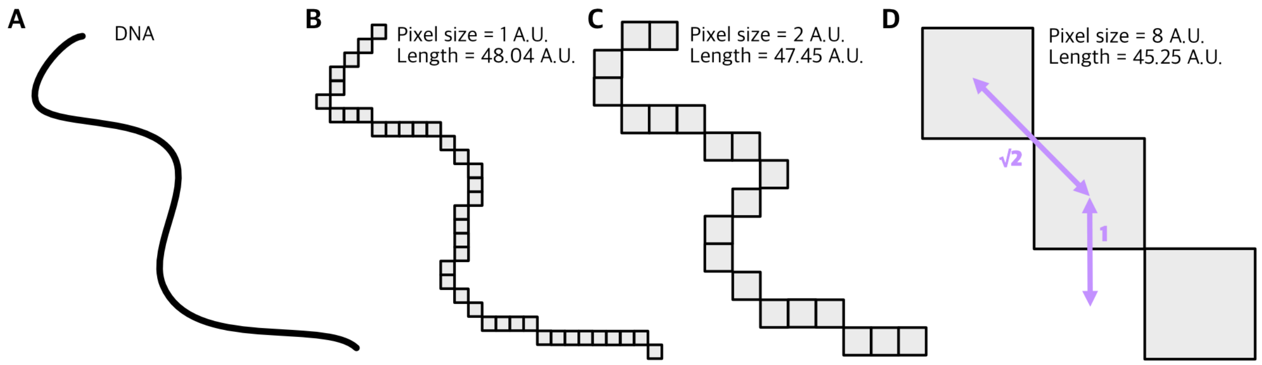


**Figure S7. The effect of pixel size on DNA length quantification.** (A) Cartoon portrayal of a DNA molecule. Cartoon portrayal of skeletonization / pixelation of DNA molecule in (A) measured with pixel sizes of (B) 1, (C) 2 and (D) 8 arbitrary units, shown only to illustrate the effect of sampling density on apparent contour length. The estimation of DNA length decreases with increasing pixel size as more details get lost with decreasing resolution.

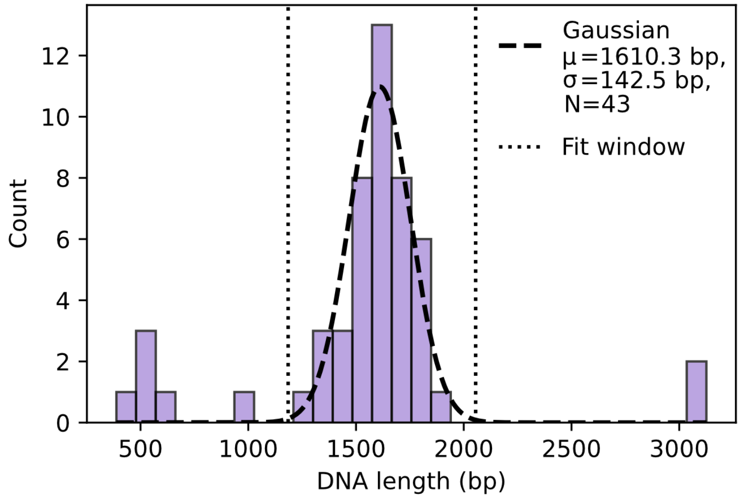


**Figure S8. Validation of DNA contour length calibration on an independent dataset.** Histogram of DNA contour lengths obtained from a single AFM image not used in the calibration procedure (Figure 2G). Gaussian fitted to the dominant peak of the distribution (black dashed line). The fit was restricted to a window centered on the histogram mode with a width determined robustly from the median absolute deviation (vertical dotted lines). Fit yielded a mean of 1610.3 bp with a standard deviation of 142.5 bp (N = 43 molecules).

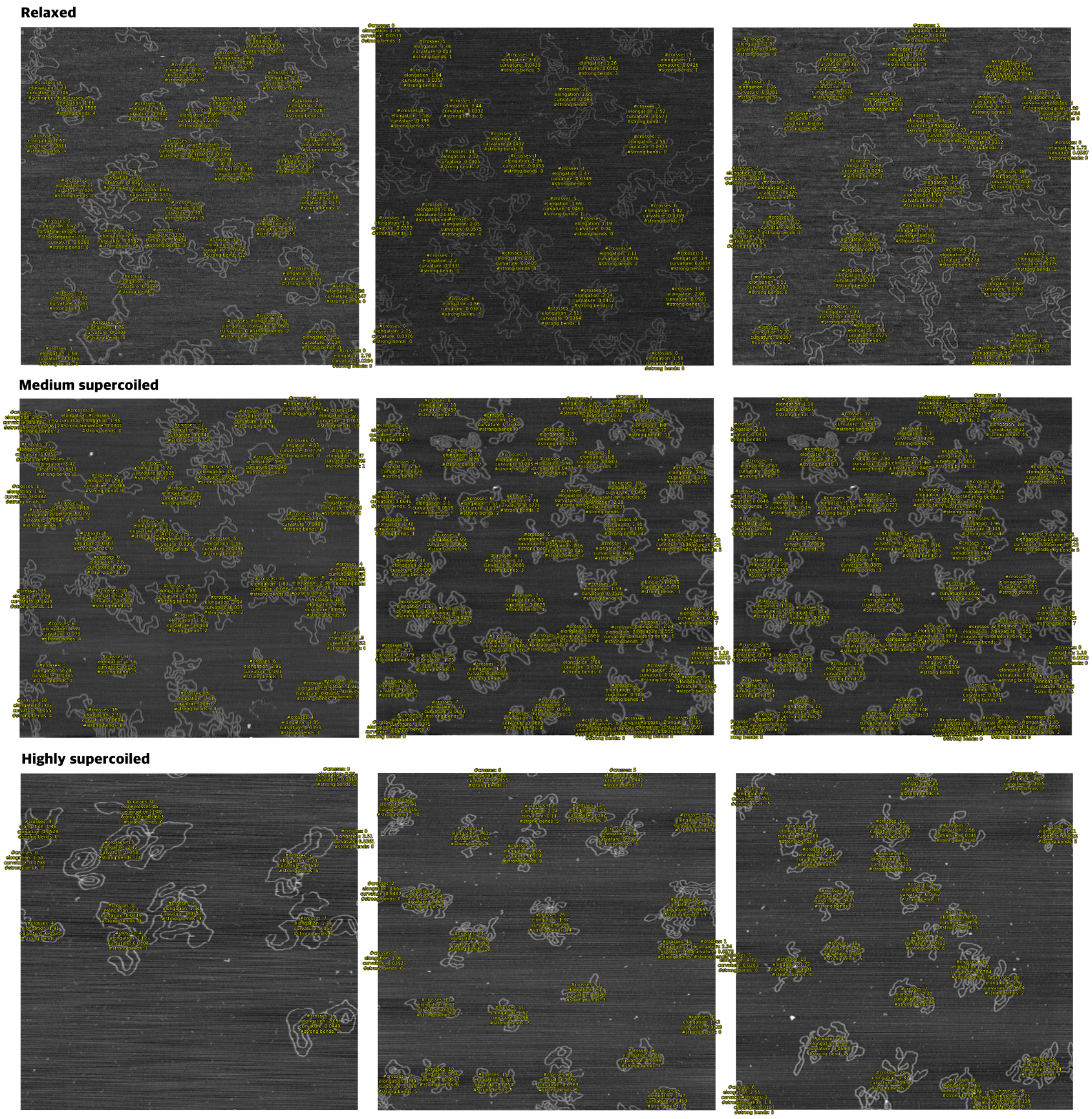


**Figure S9. Representative images with geometric features overlaid in yellow.** Displaying n.o. crosses, elongation, curvature and n.o. strong bends. Top row: relaxed DNA. Middle row: medium super coiled DNA. Bottom row: highly supercoiled DNA. Pixel size is 4 nm for all images.

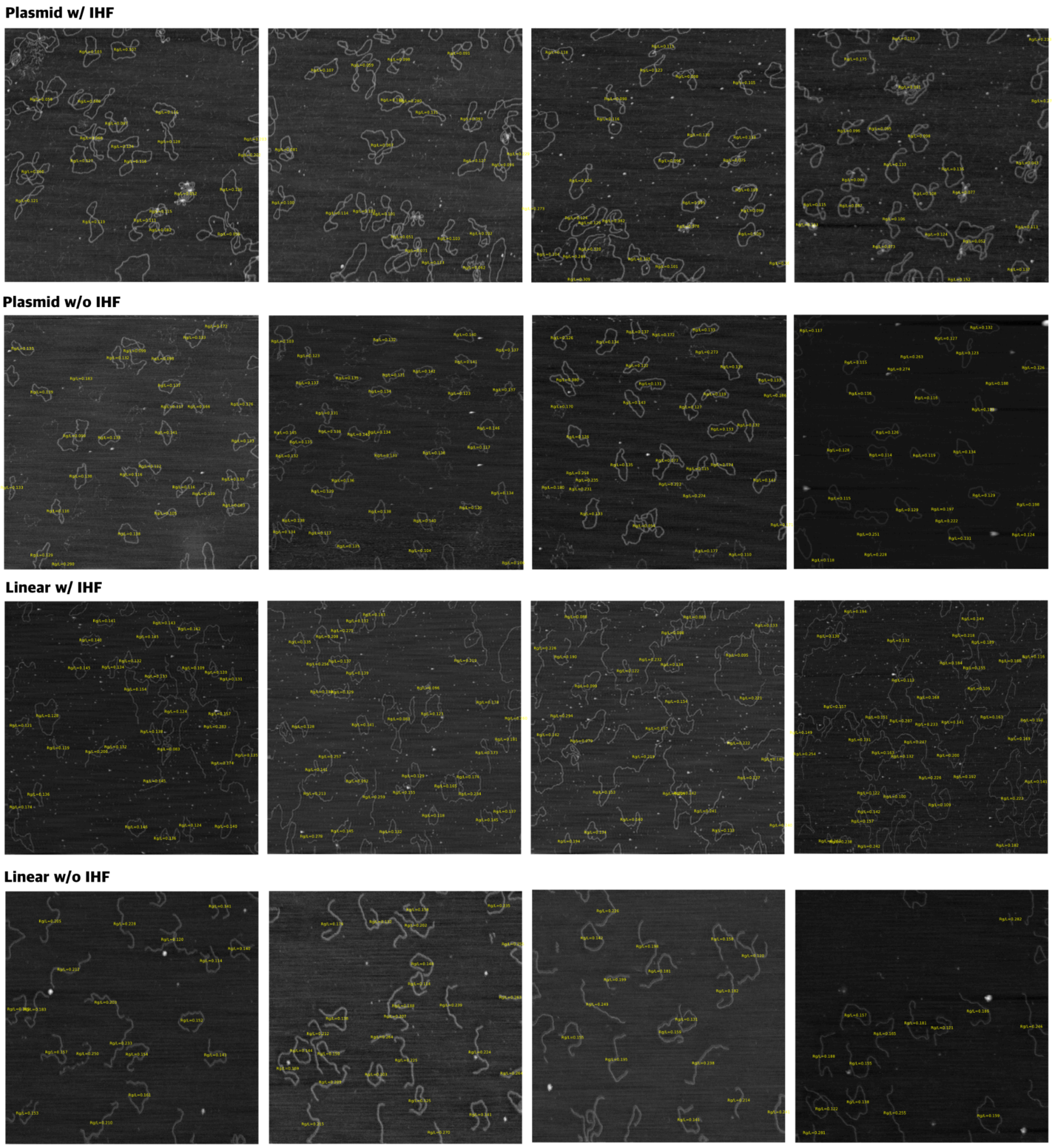


**Figure S10. Representative images of plasmid and linear DNA with and without IHF, showing compaction measure overlaid in yellow.** Top row: plasmid DNA with 50 nM IHF. Second from top row: plasmid DNA without IHF. Second from bottom row: linear DNA with 50 nM of IHF. Bottom row: linear DNA without IHF. Pixel sizes ranging from 3-4 nm.

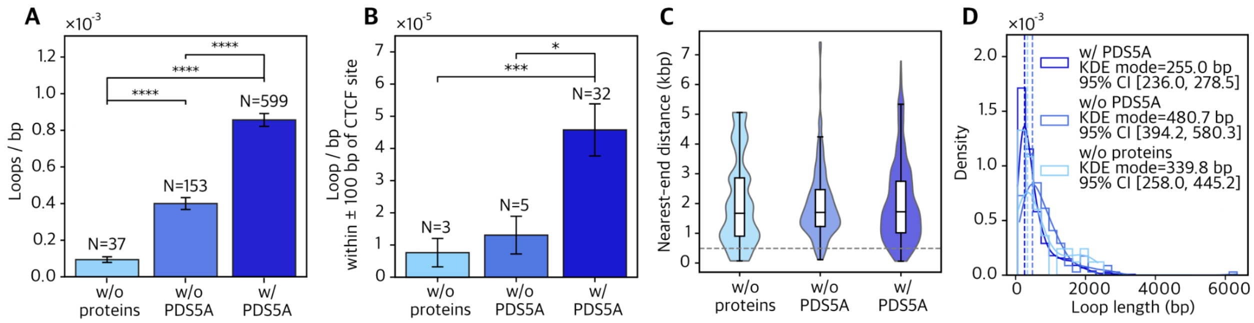


**Figure S11. Loop analysis without overlap-based filtering of highly overlapping loop candidates.** (A) Loop frequency per bp for naked DNA with no proteins present (N=37), and for DNA incubated with CTCF, cohesin-STAG1, NIPBL, and ATP in the absence (N=153) or presence (N=599) of PDS5A. (B) Loop frequency per bp within ±100 bp of the CTCF binding site for naked DNA (N=3), and for DNA incubated with proteins in the absence (N=5) or presence (N=32) of PDS5A. (C) Violin and box plots showing the nearest-end position of detected loop-like structures for naked DNA and for DNA incubated with proteins in the absence or presence of PDS5A. (D) Histograms showing the distribution of detected loop lengths in bp for naked DNA and for DNA incubated with proteins in the absence or presence of PDS5A, overlaid with kernel density estimate (KDE) fits. Dashed lines indicate the KDE mode for each condition, and the legend reports the corresponding mode estimate with its 95% confidence interval. Significance stars were assigned from two-sided Mann–Whitney U tests with Bonferroni correction for multiple comparisons. Each p-value was multiplied by the number of tested pairs, and stars were displayed according to adjusted significance thresholds: *($p < 0.05$), **($p < 0.01$), ***($p < 0.001$), and ****($p < 0.0001$); comparisons not significant after correction were left unmarked.

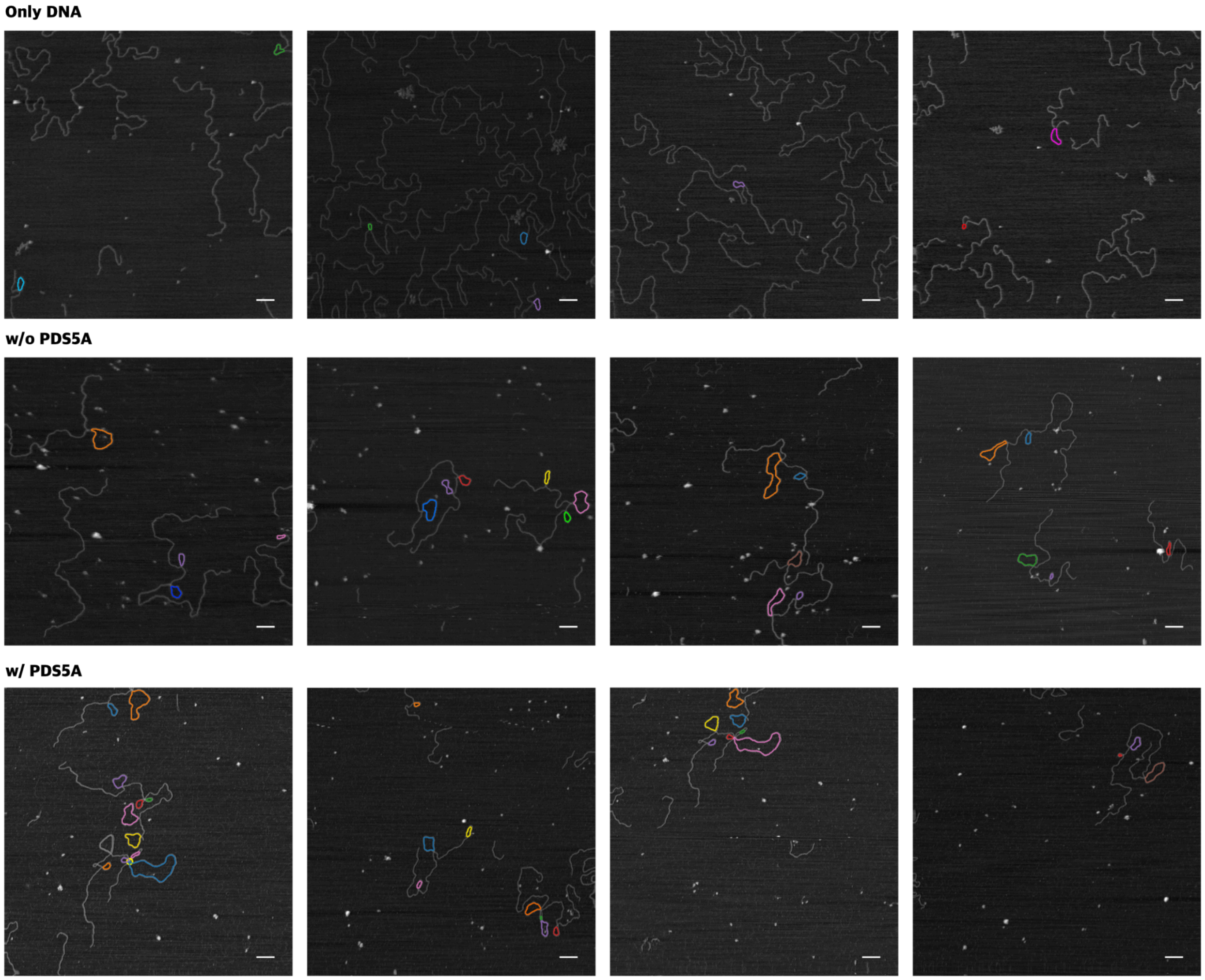


**Figure S12. Representative images with loop-like structure detection of looping experiments.** Top row: four images with loop detection overlay of no PDS5A condition. Middle row: four images with loop detection overlay of the DNA only condition, Bottom row: four images with loop detection overlay of PDS5A condition. Experimental design as detailed in main Methods and Materials. All pixel sizes are 3 nm.

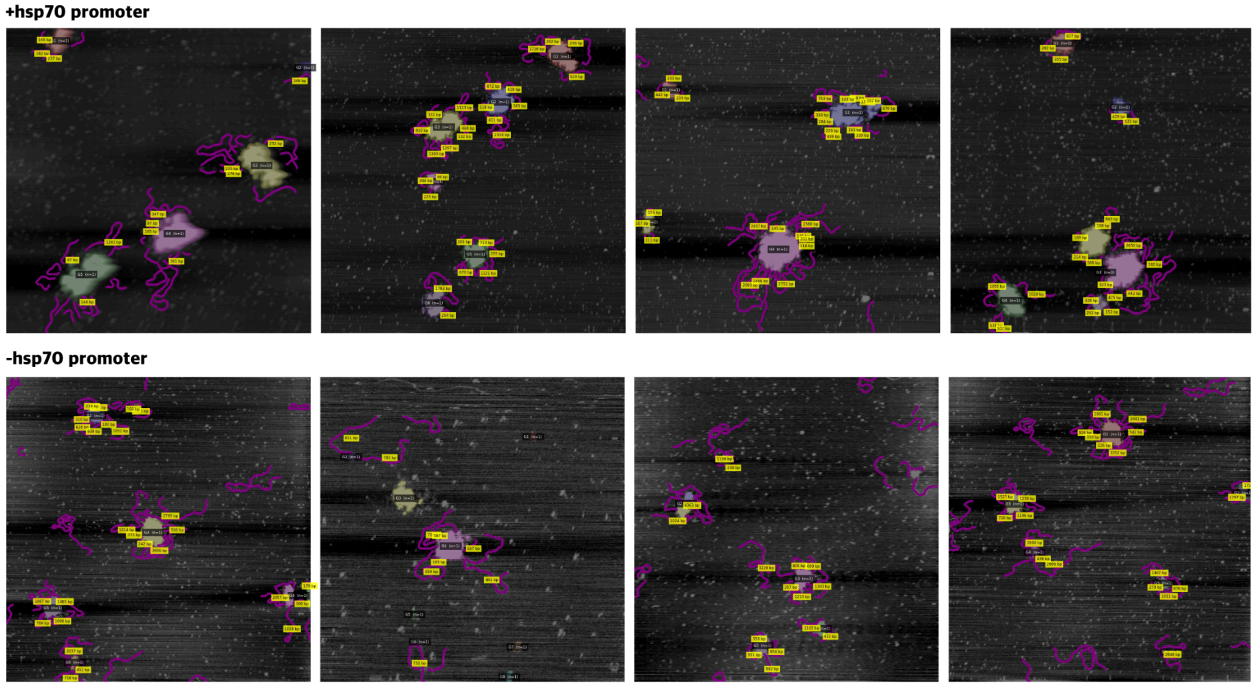


**Figure S13. Representative images of GAF clustering on linear DNA overlaid with cluster and DNA segmentation as well as their linking.** Top row: DNA with *hsp70* promoter. Bottom row: DNA without *hsp70 promoter.* All pixel sizes are 4 nm.

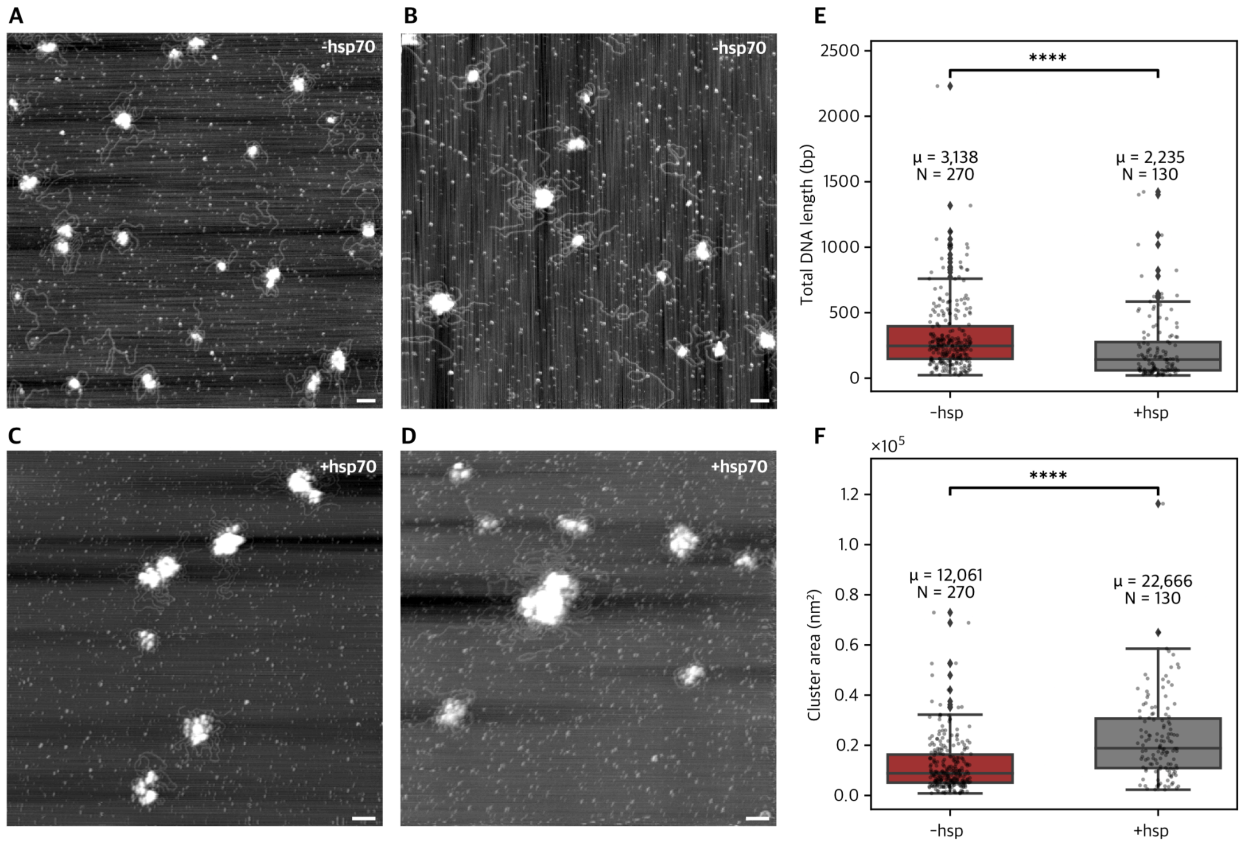


**Figure S14. Quantification of clustering and DNA recruitment of GAF with and without *hsp70* promoter.** (A/B) Representative image of GAF clustering on DNA sequence without *hsp70* promoter. (C/D) Representative image of GAF clustering on DNA sequence with *hsp70* promoter. Pixel size for all images is 4 nm. (E) Boxplots of total length of recruited DNA, to groups with a single cluster, with (mean 2235 bp, N = 130) and without (mean 3138 bp, N = 270) hsp70 promoter. (F) Boxplots of area of GAF clusters in the presence of DNA with (mean 22666 nm², N = 130) and without (mean 12061 nm², N = 270) hsp70 promoter. Clusters were filtered as detailed in main Methods and Materials.

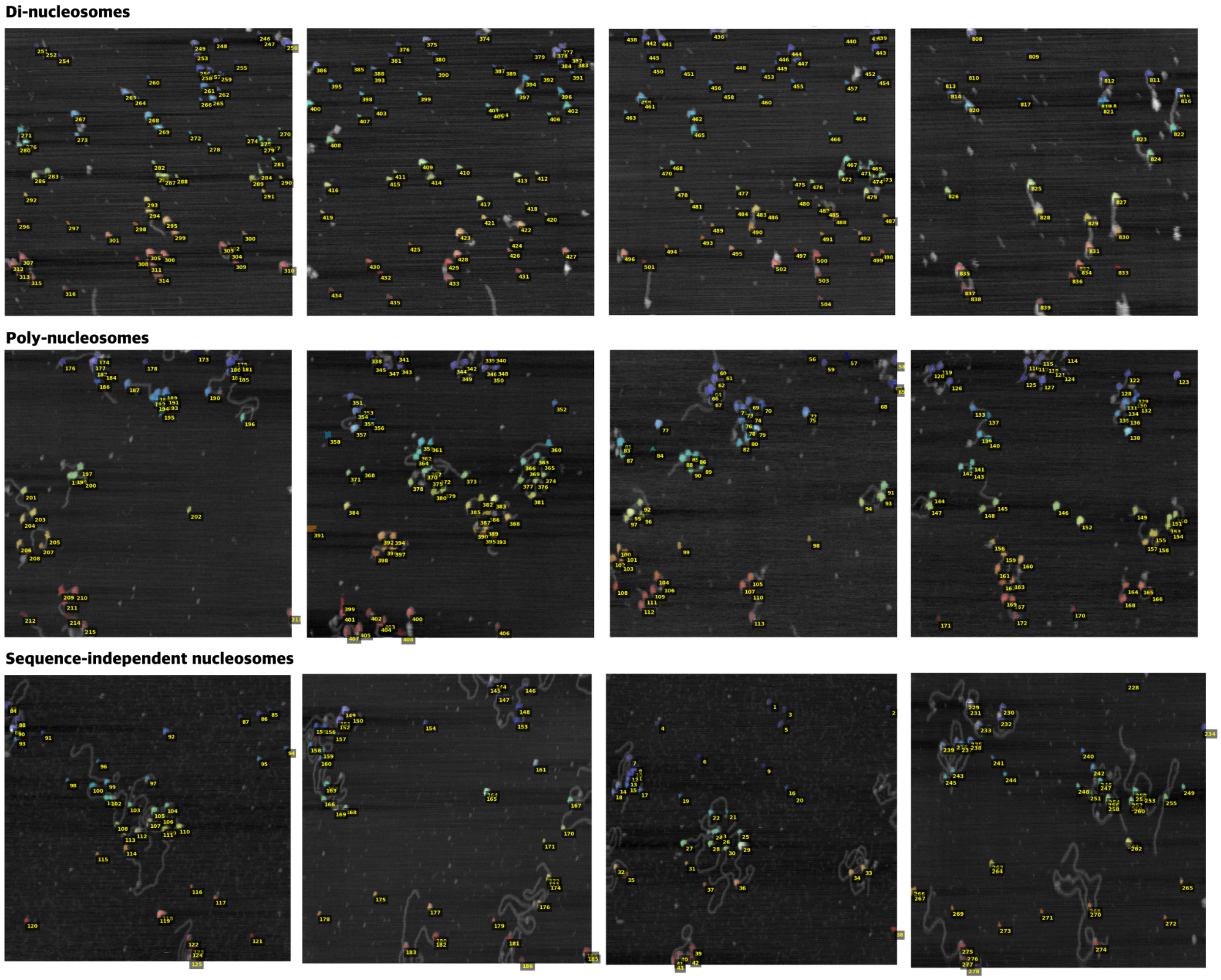


**Figure S15. Representative images and segmentation mask of nucleosome constructs.** Top row: four images with segmentation masks of the di-nucleosome construct. Middle row: four images with segmentation masks of the poly-nucleosome construct. Bottom row: four images with segmentation masks of the sequence-independent nucleosome construct. Experimental design as detailed in main Methods and Materials. Pixel sizes ranging from 1-3 nm.

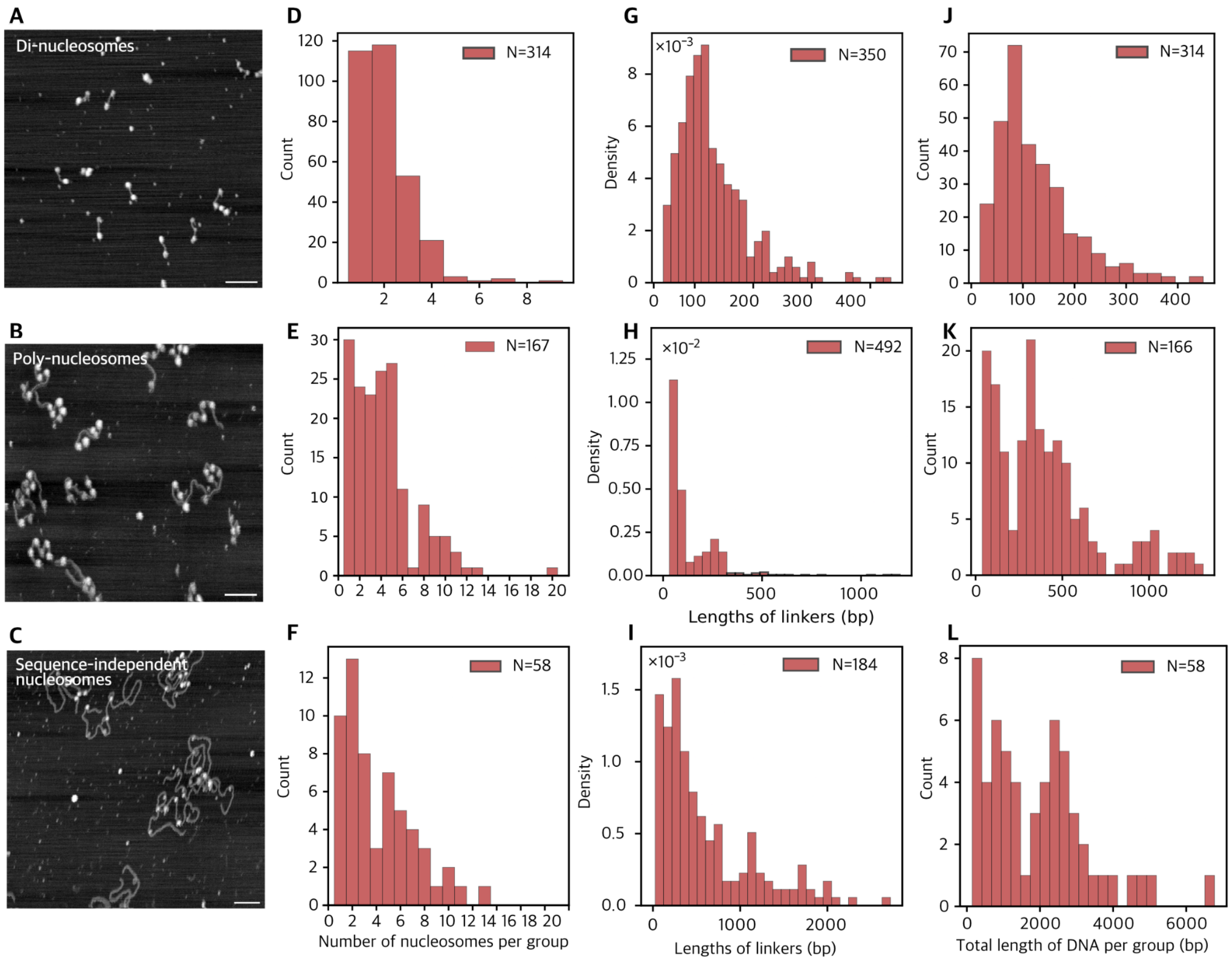


**Figure S16. Quantifications of di-nucleosomes, poly-nucleosomes and sequence-independent nucleosomes.** Representative raw image of (A) di-nuclesomes, (B) poly-nucleosomes and (C) sequence-independent nucleosomes, displayed images have pixel size 2 nm. Number of nucleosomes per group for di-nucleosomes (D), poly-nucleosomes (E) and sequence-independent nucleosomes (F). Length of nucleosome-associated DNA segments in bp for di-nucleosomes (G), poly-nucleosomes (H) and sequence-independent nucleosomes (I). Total DNA length per group in bp for di-nucleosomes (J), poly-nucleosomes (K) and sequence-independent nucleosomes (L). All distributions are unfiltered and directly output for Q4 in DNAsight.

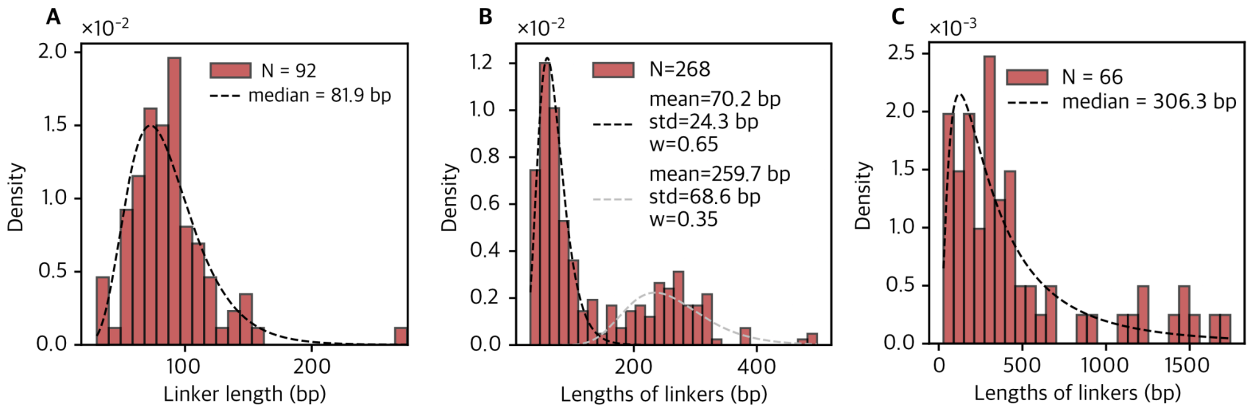


**Figure S17. Linker length distributions for DNA segments between nucleosomes.** Linker lengths were quantified only for DNA segments between nucleosomes, not including terminal DNA segments and excluding edge-touching DNA. For the (A) di-nucleosome construct, (B) hexa-nucleosome construct, and (C) sequence-independent nucleosome construct. The measured linker lengths are generally shorter than expected for the hexa-nucleosome construct. This likely reflects structural compaction of the more complex construct, which increases the probability that nucleosome clusters overlap with the DNA backbone, leading to partial obscuration and underestimation of linker lengths. Additionally, the calibration used to convert contour length to base pairs is derived from nucleosome-free DNA molecules, which are typically more extended than nucleosome-associated DNA. As nucleosome-bound DNA is more compact, this calibration could introduce a systematic underestimation of base pair lengths in these measurements.

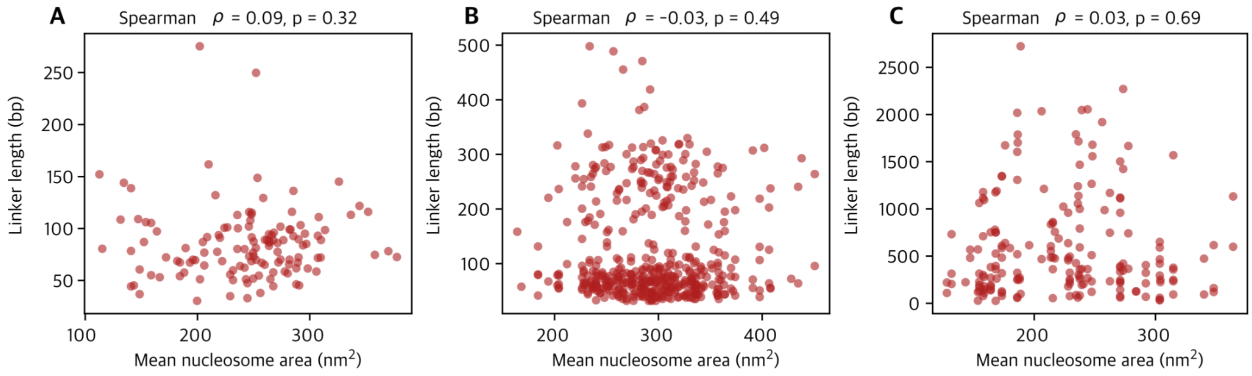


**Figure S18. Scatterplots of nucleosome-associated DNA segment lengths versus mean nucleosome size for** (A) di-nucleosome, (B) hexa-nucleosome, and (C) sequence-independent nucleosome constructs. Spearman's rank correlations were weak and not statistically significant in all three cases: di-nucleosome, $\rho=0.09$, $p=0.32$ hexa-nucleosome, $\rho=-0.03$, $p=0.49$; sequence-independent nucleosomes, $\rho=0.03$, $p=0.69$.
